# Higher-Order Organization Links Circulating Metabolites to Glucocorticoid Action in Human Adipose Tissue

**DOI:** 10.64898/2026.10.08.757658

**Authors:** Kate Bennett, Dimitrios Chantzichristos, Terence Garner, Per-Anders Jansson, Anja Meyer, Eleanor Doman, Johanna McQueen, Per-Arne Svensson, Oskar Ragnarsson, Gudmundur Johannsson, Adam Stevens

## Abstract

Glucocorticoids (GCs) are regulators of metabolism, yet how physiological GC exposure remodels adipose metabolism and its relationship with circulating biomarkers remains poorly understood.

In a randomised cross-over study of individuals with primary adrenal insufficiency, near-physiological GC exposure was compared to withdrawal using integrated serum metabolomics, abdominal subcutaneous microdialysis metabolomics, and adipose tissue transcriptomics.

GC exposure induced coordinated changes in adipose lipid and amino-acid metabolism. Pathway-level entropy analysis revealed GC-state-dependent transcriptional reorganisation, with concentrated organisation during exposure and diffuse organisation during withdrawal.

Integrative hypergraph and topological analyses revealed GC-state-dependent reorganisation of circulating metabolite-adipose tissue relationships, with greater higher-order coordination during exposure. The circulating phenylalanine/kynurenine ratio discriminated GC state, showed the strongest shared connectivity to GC-regulated adipose networks, persisted at midnight and in an independent dose-response study, and exhibited circadian rhythmicity.

These findings reveal dynamic GC regulation of adipose metabolism and identify a circulating biomarker with potential to guide personalised GC therapy

## INTRODUCTION

Glucocorticoids (GCs) are steroid hormones essential for human life and health, regulating immune, metabolic, and central nervous system function^1^. GCs are widely used to treat inflammatory disorders, as part of cancer therapy, and as lifelong life-saving replacement therapy in patients with adrenal insufficiency^2,3^. Long-term GC treatment is associated with adverse metabolic and cardiovascular effects^4^, and increased mortality^5,6^. Despite their widespread use, approaches to individualise dose titration and optimise efficacy and safety are limited and fail to capture tissue-specific GC action.

Adipose tissue is a key target of GC action and plays a central role in mediating their metabolic effects^7^. Alongside liver and muscle, adipose tissue coordinates energy mobilisation in response to the increased GC exposure that occurs during stress, yet GC signalling in adipocytes is complex and context dependent. GCs can promote pre-adipocyte differentiation, impair insulin-stimulated glucose uptake, alter lipid handling, and induce lipolysis in mature adipocytes^7-10^. Chronic GC excess, as observed in Cushing’s syndrome or during prolonged pharmacological GC therapy, is associated with a shift towards central/visceral fat accumulation through mechanisms that are not yet fully understood^7^.

In humans, the main endogenous GC, cortisol, is secreted in a pronounced circadian rhythm, with a morning peak and nocturnal nadir. This rhythmic exposure regulates both central and peripheral clocks, and disruption of this circadian pattern has adverse consequences for metabolic health^11^. Moreover, adipocyte metabolism shows daily variation in both animals and humans^12,13^, and experimental flattening of the GC circadian rhythm promotes adipocyte hypertrophy and fat accumulation independent of total GC exposure^14^. Thus, GC action in adipose tissue is shaped not only by total exposure, but also by circadian timing, tissue context, and downstream metabolic state.

GC signalling within tissues is shaped by pre-receptor metabolism, receptor-mediated interactions with transcriptional networks, non-genomic pathways, and the temporal pattern of exposure^4,11,15^. These mechanisms also regulate the tissue-specific actions of GCs, which altogether contribute to the global regulation of GC action on metabolism and immune function. Due to this complexity, current clinical and biochemical measures of GC exposure can only identify gross GC excess or deficiency and lack the resolution to capture physiologically appropriate GC action.

Together, these considerations suggest that GC action is best understood not as isolated changes in individual biomarkers, but as coordinated regulation of multi-pathway responses that vary across tissues and circadian phase. We therefore hypothesised that circulating markers of adipose tissue GC action would be better identified using analytical approaches that move beyond single-molecule readouts and pairwise associations, to capture higher-order metabolic reorganisation and cross-tissue coordination.

To address these challenges, we used a human experimental model of primary adrenal insufficiency, in which endogenous cortisol production is absent, to investigate adipose tissue responses under tightly controlled conditions of near-physiological GC exposure and withdrawal. By combining abdominal subcutaneous microdialysis metabolomics with adipose tissue transcriptomic profiling, and integrating these data with serum metabolomics, we aimed to define adipose tissue GC metabolic action and determine whether circulating metabolites can serve as biomarkers of downstream adipose tissue-specific GC responses. A graphical overview of the study is presented in Fig. 1.

**Figure 1.**
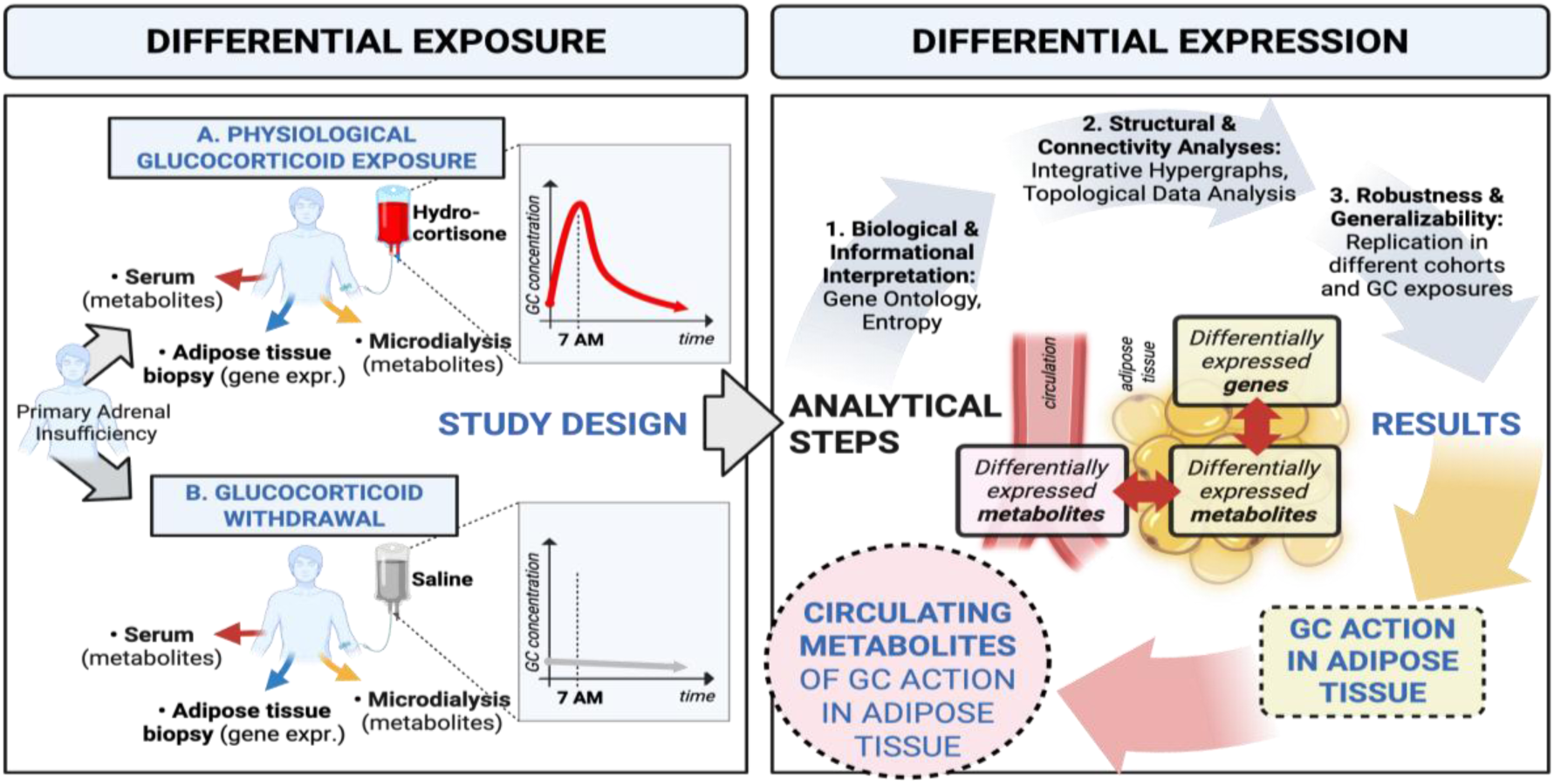
Graphical Overview. Randomised, single-blinded, cross-over study (BIOCORT) in patients with primary adrenal insufficiency (Addison’s disease), who completely lack endogenous glucocorticoid (GC) production. The left panel (A–B) depicts the differential exposure conditions—near physiological GC replacement with hydrocortisone infusion versus GC withdrawal with saline, and sampling at 07:00, corresponding to the normal GC peak in healthy individuals. At each condition, fasting blood, abdominal subcutaneous dialysates (microdialysis) and abdominal subcutaneous adipose tissue (biopsy), were collected for metabolomic and transcriptomic profiling. The right panel summarises the analytical workflow of the differential expression, including multi-layer integration of gene and metabolite changes, structural and connectivity analyses using hypergraph-based models, and biological interpretation through pathway and ontology analyses. Putative metabolic markers in the circulation reflecting GC action in adipose tissue were validated in midnight samples and independent study groups with distinct GC exposure patterns. Together, this design enables mechanistic studies of GC in adipose tissue under physiological conditions together with identification of circulating markers of GC action in adipose tissue. Figure created using BioRender.com.

## RESULTS

To define how GCs regulate human adipose tissue and how these effects are reflected in the circulation, we analysed metabolic and transcriptomic responses under controlled GC exposure and withdrawal (Fig. 1). We first confirmed effective modulation of GC signalling and minimal confounding physiological changes. We then characterised GC-dependent changes in adipose tissue metabolism and transcriptomic network organisation, before examining whether adipose metabolic responses were mirrored in the circulation. Next, we applied systems-level higher-order network analysis to define cross-tissue coordination. Finally, we identified circulating metabolite markers that capture adipose tissue GC action and assessed their reproducibility in independent studies with different GC-exposure datasets.

### GC exposure remodels adipose tissue metabolism

To investigate GC-dependent metabolic changes in human adipose tissue, we studied ten individuals with primary adrenal insufficiency, Addison’s disease, undergoing controlled near-physiological GC exposure, and GC withdrawal in the BIOCORT study^16^. Hydrocortisone infusion achieved physiological morning serum cortisol concentrations, whereas concentrations remained low during saline infusion, confirming clear separation between GC states at both the morning peak and midnight trough. Metabolic and clinical parameters remained stable between interventions, except for a modest increase in cystatin C during exposure, indicating minimal confounding physiological perturbations (Extended Data Table 1). Expression of known GC-responsive genes in adipose tissue (e.g. *FKBP4, DCK)* was increased during GC exposure, confirming tissue-level activation of GC signalling^17^.

**Extended Data Table 1.**
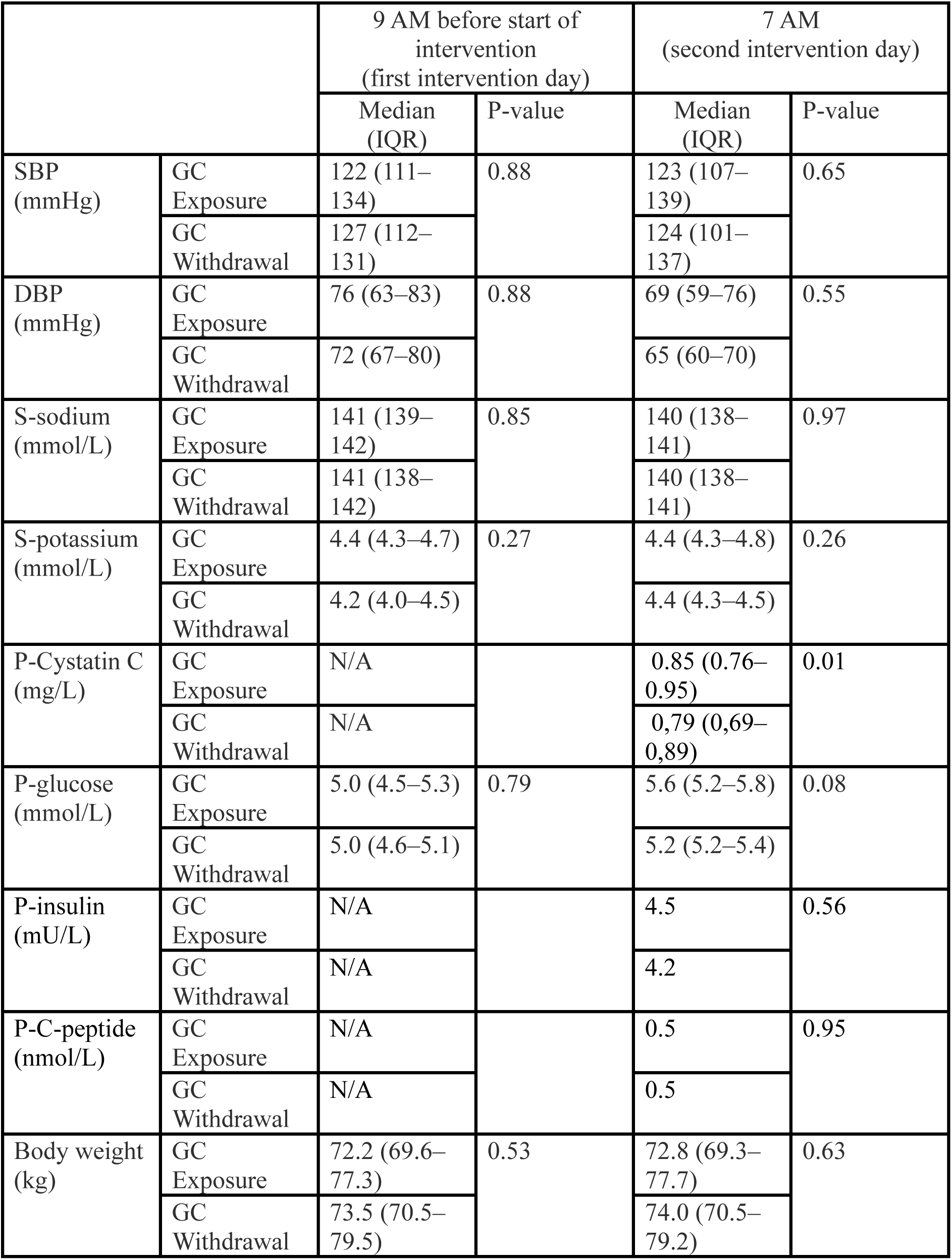
Selected anthropometric, clinical, and biochemical outcomes measured before the start of interventions and at 07:00 during both study arms, with a focus on variables related to glucose and lipid metabolism. Adapted from previous publications^16,17^. Abbreviations: S, serum; P, plasma; SBP, systolic blood pressure; DBP, diastolic blood pressure; N/A, not applicable; IQR, interquartile range.

### GC exposure induces a discrete metabolic signature in human adipose tissue

To determine whether near-physiological GC exposure alters adipose tissue metabolism, we profiled metabolites in abdominal subcutaneous adipose interstitial fluid using in situ microdialysis. This approach samples the local extracellular metabolic environment in vivo, providing a dynamic tissue-level readout that is distinct from both circulating metabolite profiles and bulk adipose tissue measurements. To assess potential confounders, we examined associations of biochemical, clinical and adipose transcriptomic variables with principal components (PCs). Total GC exposure (cortisol AUC) aligned most strongly with the first principal component (PC1), which captures the largest proportion of the overall variance (Fig. 2a). Other biological variables, including markers of insulin sensitivity (expression of *ADIPOQ* and *SLC2A4*), and inflammation (expression of *CD68*)^18-20^ showed weaker associations with the major components. BMI showed the largest observed association with PC5, which explained only 6.6% of the variance. Supervised multivariate analysis demonstrated separation of adipose metabolic profiles between GC exposure and withdrawal (Fig. 2b).

**Figure 2.**
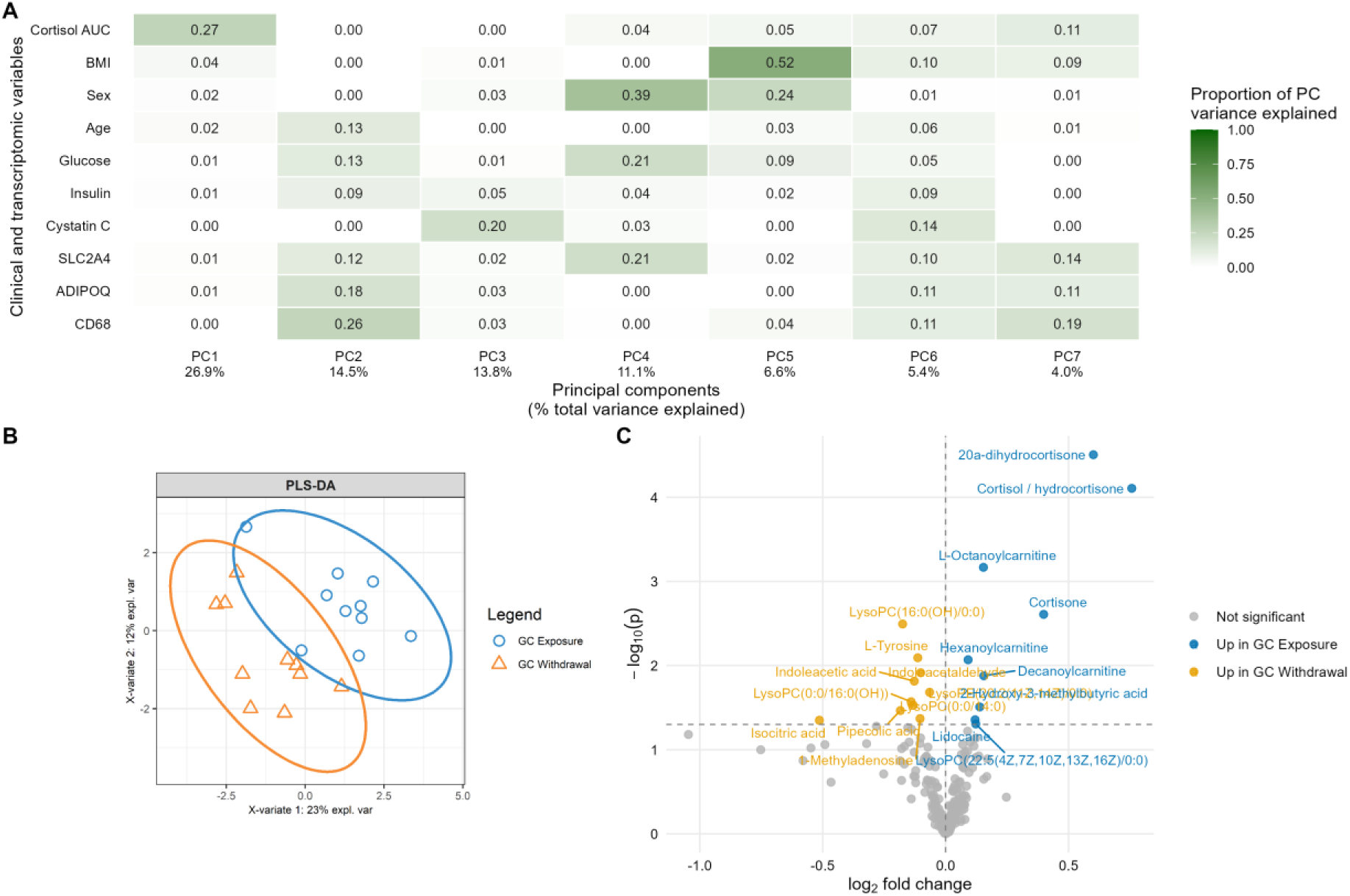
Adipose tissue metabolomics. **(A)** Heatmap showing the proportion of variance in each adipose tissue metabolomic principal component (PC) explained by selected clinical variables and adipose tissue gene-expression markers related to insulin sensitivity (ADIPOQ and SLC2A4) and inflammation (CD68). The percentage of total metabolomic variance explained by each principal component is shown beneath the x-axis. (**B)** Partial least squares discriminant analysis (PLS-DA) illustrating separation of adipose tissue metabolomic profiles between GC exposure (blue circles) and GC withdrawal (orange triangles). (**C)** Volcano plot showing metabolites differing between GC exposure and withdrawal. Blue points on the right indicate metabolites increased during GC exposure, orange points on the left indicate metabolites decreased during GC exposure, and grey points indicate non-significant metabolites.

Differential abundance analysis identified 19 adipose tissue metabolites that differed between GC exposure and GC withdrawal (paired t-test P-value < 0.05) (Fig. 2C; Extended Data Table 2). Two steroid metabolites, 20α-dihydrocortisone and cortisol/hydrocortisone, remained significant following Benjamini-Hochberg correction (FDR < 0.05). The remaining metabolites are presented as exploratory findings.

Cortisol and cortisone derivatives were increased during GC exposure, confirming local GC availability within adipose tissue. GC exposure was associated with coordinated changes across lipid, amino acid, nucleoside and organic-acid pathways.

Several medium-chain acylcarnitines were increased during GC exposure, consistent with altered fatty-acid handling and changes in lipid mobilisation and mitochondrial fatty-acid oxidation. Isocitric acid was decreased during GC exposure, consistent with reports on altered TCA-cycle activity^21^. Several glycerophospholipid species were also altered, with both increases and decreases suggesting species-specific phospholipid remodelling under physiological GC exposure.

GC exposure was also associated with changes in amino-acid-related metabolites. Tyrosine, pipecolic acid, indoleacetaldehyde, and indoleacetic acid were lower during GC exposure, indicating altered aromatic amino acid and tryptophan-related metabolism within adipose tissue. In contrast, GC exposure increased 2-hydroxy-3-methyl butyric acid, a valine-derived metabolite, suggesting altered branched-chain amino acid metabolism. Finally, 1-methyladenosine was reduced during GC exposure, suggesting altered nucleoside metabolism as an additional component of the adipose tissue GC response. Together, these data identify a distinct adipose metabolic response to physiological GC exposure.

**Extended Data Table 2.** Differentially abundant metabolites in adipose tissue microdialysate between GC exposure and GC withdrawal. Paired t-test between GC exposure and withdrawal adipose tissue metabolomics using MetaboAnalyst. Showing log2 fold change (GC exposure relative to GC withdrawal), P-value, FDR-adjusted P-value, and HMDB class. Lidocaine, an exogenous metabolite, was administered subcutaneously prior to sampling during both interventions.

| Metabolite | log2FC | P-value | FDR | HMDB Class |
| --- | --- | --- | --- | --- |
| 20α-dihydrocortisone | 0.601 | 3.110E-05 | 0.007 | Steroid and Steroid Derivatives |
| Cortisol / hydrocortisone | 0.757 | 7.805E-05 | 0.009 | Steroid and Steroid Derivatives |
| L-Octanoylcarnitine | 0.154 | 0.001 | 0.052 | Fatty Acyls |
| Cortisone | 0.399 | 0.002 | 0.140 | Steroid and Steroid Derivatives |
| LysoPC(16:0(OH)/0:0) | -0.175 | 0.003 | 0.146 | Glycerophospholipid |
| L-Tyrosine | -0.113 | 0.008 | 0.278 | Carboxylic acids and derivatives |
| Hexanoylcarnitine | 0.091 | 0.009 | 0.278 | Fatty Acyls |
| Indoleacetaldehyde | -0.101 | 0.012 | 0.336 | Indoles and Derivatives |
| Decanoylcarnitine | 0.156 | 0.013 | 0.336 | Fatty Acyls |
| Indoleacetic acid | -0.128 | 0.015 | 0.350 | Indoles and Derivatives |
| LysoPE(20:2(11Z,14Z)/0:0) | -0.065 | 0.021 | 0.432 | Glycerophospholipid |
| LysoPC(0:0/16:0(OH)) | -0.140 | 0.027 | 0.507 | Glycerophospholipid |
| LysoPC(0:0/14:0) | -0.132 | 0.030 | 0.507 | Glycerophospholipid |
| 2-Hydroxy-3-methylbutyric acid | 0.139 | 0.031 | 0.507 | Fatty Acyls |
| Pipecolic acid | -0.184 | 0.034 | 0.522 | Carboxylic acid and derivatives |
| 1-Methyladenosine | -0.104 | 0.043 | 0.566 | Purine Nucleoside |
| Lidocaine | 0.120 | 0.044 | 0.566 | Benzene and substituted derivatives |
| Isocitric acid | -0.514 | 0.045 | 0.566 | Carboxylic acid and derivatives |
| LysoPC(22:5(4Z,7Z,10Z,13Z,16Z)/0:0) | 0.122 | 0.049 | 0.572 | Glycerophospholipid |

### GC exposure remodels adipose metabolic pathway structure at the transcriptomic level

To determine whether these observed metabolic changes were accompanied by broader reorganisation of adipose tissue metabolism, we analysed pathway-level transcriptomic structure using hypergraph-based network representations. Conventional networks capture pairwise relationships between genes, where edges represent direct associations between pairs of genes. Hypergraphs extend this framework by allowing a single hyperedge to connect multiple nodes simultaneously, thereby representing higher-order relationships among groups of genes. This may better model complex biological systems, as genes often act within multi-component processes rather than as isolated pairwise interactions.

Within this higher-order framework, we quantified pathway organisation using entropy, which captures how shared higher-order interactions are distributed across gene pairs within a pathway. In this context, low entropy indicates that shared interactions are concentrated among a subset of gene pairs, and higher entropy reflects more even distribution of connectivity. In this analysis, we assessed whether GC exposure altered the organisation of metabolic pathways at the transcriptomic level.

Across all metabolic pathways examined, entropy differed between GC exposure and withdrawal (Fig. 3a), with this difference larger than size-matched random gene sets, indicating pathway-specific reorganisation beyond global transcriptome-wide effects (Fig. 3b). Most pathways, including lipid biosynthesis and catabolism, exhibited lower entropy during GC exposure, indicating that shared interactions were concentrated among specific gene pairs. In contrast, GC withdrawal was associated with higher entropy across these pathways, reflecting a redistribution of shared interactions.

**Figure 3.**
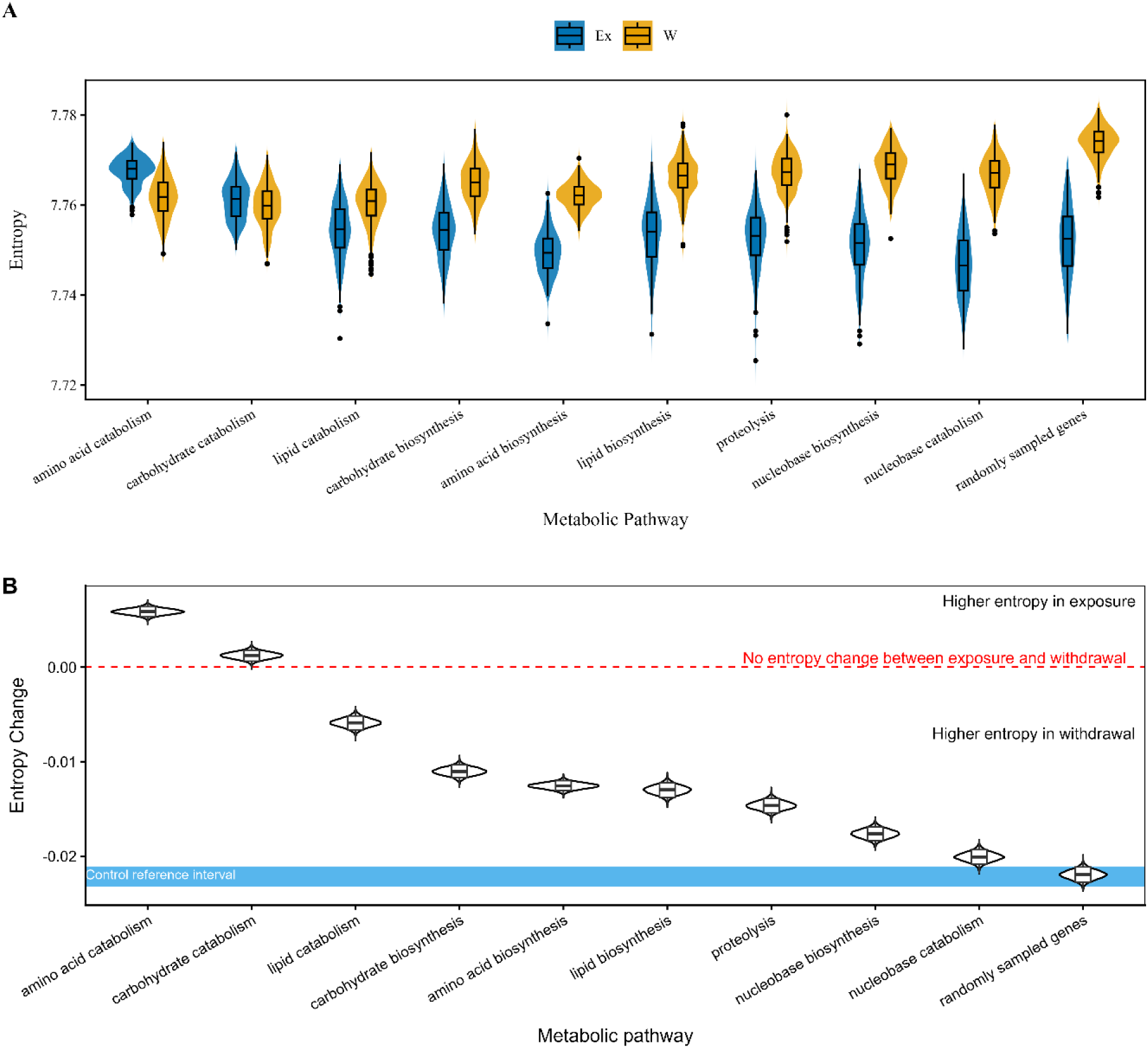
Entropy changes in adipose tissue metabolic pathways between GC exposure and withdrawal. **A)** Pathway-specific entropy of adipose tissue transcriptomic hypergraph networks during GC exposure (blue) and GC withdrawal (orange). For each metabolic pathway, nodes represent genes within that pathway, and hyperedges represent shared associations with the rest of the adipose transcriptome, capturing pathway-level organisation beyond pairwise relationships. High entropy indicates a more even distribution of connections across the network, while low entropy indicates a more clustered organisation, with connections concentrated among a subset of genes. **B)** Bayesian comparison of entropy differences between GC states. Entropy change is shown as the difference between GC exposure and GC withdrawal, meaning positive values indicate higher entropy during GC exposure, whereas negative values indicate higher entropy during GC withdrawal. The blue shaded band represents the distribution of entropy changes obtained from randomly sampled genes, serving as a transcriptome-wide negative control. The red dashed line marks no difference in entropy between GC states.

Notably, carbohydrate and amino acid catabolic pathways displayed the opposite pattern, with increased entropy during GC exposure. This suggests redistribution of metabolic control rather than uniform suppression or activation. These findings indicate that GC exposure does not simply shift metabolic activity uniformly, but instead alters the organisation of metabolic pathway-associated transcriptomic networks within adipose tissue.

Collectively, adipose metabolic and transcriptomic analyses demonstrate that physiological GC exposure drives a distinct and coordinated metabolic state within human adipose tissue. GC exposure reorganises lipid and amino acid metabolism through pathway-specific changes in metabolite abundance and transcriptional organisation. In contrast, GC withdrawal was associated with a more distributed interaction architecture, indicating that both GC peaks and troughs actively shape the circadian adipose tissue metabolic regulation.

### GC exposure reorganises coordinated metabolic signals between adipose tissue and the circulation

#### GC-dependent changes in adipose-circulation metabolic coupling

To determine whether GC-induced changes in adipose tissue metabolism were reflected in the circulating metabolome, we examined the relationship between matched metabolites in the adipose interstitial fluid and serum. Correlation analysis revealed widespread concordance between adipose and circulating metabolites under both GC exposure and withdrawal (Fig. 4a). However, a subset of metabolites, including kynurenine, proline, and selected acylcarnitines, exhibited GC-dependent changes in the strength or direction of their adipose-serum correlations. This indicates that GC exposure not only alters metabolite abundance, but also changes how closely circulating metabolite levels reflect local adipose tissue metabolism. Thus, GC signalling reorganises the relationship between adipose tissue and the circulation, beyond static changes in metabolite concentrations. However, these pairwise relationships provide only a partial view of cross-tissue organisation. Given the pathway-specific restructuring observed within adipose tissue, we next asked whether GC exposure similarly alters the higher-order organisation linking adipose tissue and circulating metabolites. Addressing this question required approaches that move beyond pairwise correlations to capture coordinated relationships among groups of molecular features (Fig. 4b).

**Figure 4.**
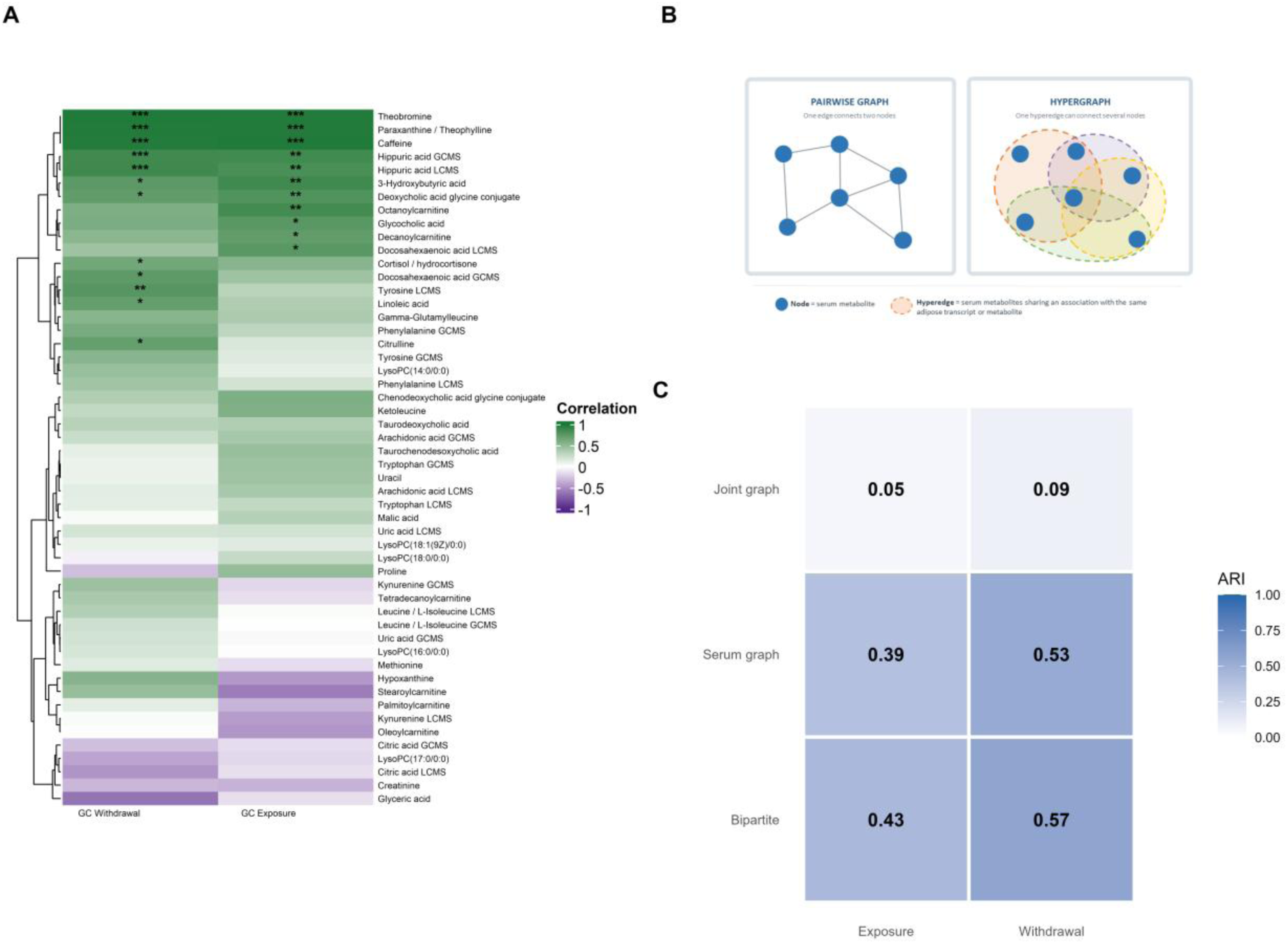
Comparison of metabolite associations and network-derived module structures during GC exposure and withdrawal. **A)** Heatmap showing correlation values for matched metabolites present in the circulation and adipose tissue. Metabolites were correlated in both GC exposure and GC withdrawal, with hierarchical clustering. Green depicts positively correlated metabolites, while purple shows those negatively correlated. Asterisks indicate statistically significant associations (p < 0.05, p < 0.01, p < 0.001). **B)** Schematic illustration of the network representations used for module detection. In the pairwise graph, edges connect pairs of serum metabolites. In the hypergraph, hyperedges connect groups of serum metabolites sharing an association with the same adipose transcript or metabolite. **(C)** Agreement between modules identified from the hypergraph and alternative network representations, quantified using the Adjusted Rand Index (ARI). Comparisons were performed against a serum-only correlation network, a joint pairwise network including both serum metabolites and adipose omics features, and a bipartite network preserving direct serum-adipose associations. Networks for both GC exposure and withdrawal conditions were constructed from the same underlying serum metabolite and adipose omics datasets but differed in how relationships were represented. Higher ARI values indicate greater similarity between module assignments.

#### Hypergraphs capture non-random organisation beyond pairwise networks

To characterise this higher-order organisation, we integrated serum metabolomics with adipose tissue transcriptomic and metabolic data using hypergraph-based network models. This approach enabled us to examine how groups of serum metabolites collectively relate to metabolites and transcripts within adipose tissue, thereby characterising systems-level metabolic organisation across tissues.

To determine whether the hypergraph representation captured structure beyond conventional pairwise networks, we compared hypergraph-derived serum metabolite modules with modules obtained from alternative network representations. Hypergraph-derived modules most closely resembled those from the bipartite representation, which preserves direct serum–adipose associations, and showed moderate agreement with serum-only modules (Fig. 4c; Supplementary Fig. 1a). Agreement was much lower with the joint pairwise graph. Thus, hypergraph organisation retained information arising from direct cross-tissue relationships while producing higher-order groupings not recovered when all features were combined in a conventional pairwise network. Null-model testing supported non-random organisation, with relationships exceeding chance expectations (FDR-adjusted q < 0.05; Supplementary Fig. 1b-c). We therefore used the hypergraph as the primary integrative framework to quantify shared connectivity between circulating metabolites and adipose multi-omic features, and to evaluate candidate biomarkers within GC-responsive adipose networks.

#### GC state alters the organisation of adipose-serum hypergraph connectivity

Having established that the hypergraph captured complementary organisation not fully reproduced by pairwise network representations, we next compared its connectivity structure between GC states. Overall density and hyperedge-size profiles were similar between exposure and withdrawal (Extended Data Fig. 1 a,b). In contrast, exposure showed a broader node hyperdegree distribution than withdrawal, indicating greater heterogeneity in the number of adipose features associated with individual serum metabolites (Fig. 5a). Exposure also showed a right-shifted distribution of shared connectivity, calculated as the summed co-degree of each serum metabolite with all other serum metabolites (Fig. 5b). Thus, exposure was characterised by more uneven direct serum-adipose connectivity and greater sharing of adipose-linked associations among circulating metabolites.

**Figure 5:**
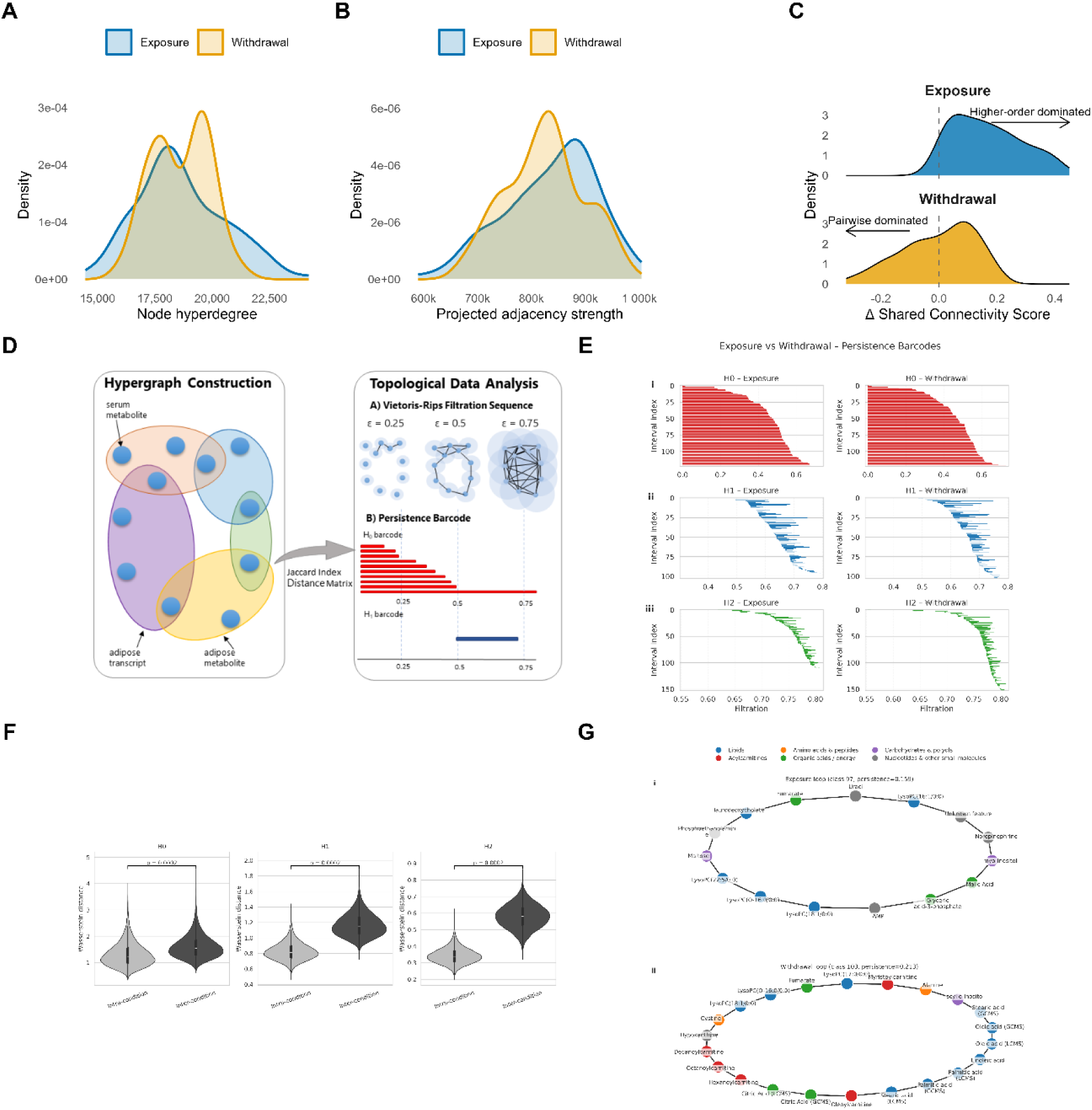
Hypergraph topology and connectivity in integrated serum metabolite-adipose feature networks across GC exposure and withdrawal. **(A)** Distribution of node hyperdegree across serum metabolites in GC exposure and withdrawal hypergraphs. Node hyperdegree represents the number of adipose transcriptomic and metabolic features associated with each serum metabolite and therefore quantifies direct serum-adipose incidence connectivity. **(B)** Distribution of shared connectivity across serum metabolites. Shared connectivity was calculated for each serum metabolite as its summed co-degree with all other serum metabolites, where pairwise co-degree represents the number of adipose features jointly associated with a pair of serum metabolites. **(C)** Relative contribution of shared connectivity compared with direct incidence connectivity. Node hyperdegree and shared connectivity were min-max normalised separately within each GC state, and the Δ shared-connectivity score was calculated as normalised summed co-degree minus normalised node hyperdegree. Right-shifted distributions indicate greater contribution of higher-order interactions, whereas left-shifted distributions indicate dominance of independent pairwise associations to adipose features. **(D)** Workflow for integrating serum metabolomics with adipose tissue metabolomics and transcriptomics to generate hypergraphs, followed by conversion to distance matrices and topological data analysis using Vietoris-Rips filtration and persistence barcode computation. **(E)** Persistence barcodes reveal topological differences between GC exposure and withdrawal hypergraphs. Barcodes are shown for H₀ (connected components), H₁ (loops), and H₂ (voids). **(F)** Wasserstein distances between persistence diagrams across homology dimensions. Violin plots show intra-group (exposure-exposure, withdrawal-withdrawal) and inter-group (exposure-withdrawal) distances calculated from hypergraphs generated by subsampling 80% of serum metabolite nodes without replacement. Greater inter-group distances indicate increased topological dissimilarity between exposure and withdrawal states. Statistical significance was assessed using label-based permutation tests (5,000 permutations; Extended Data Fig. 1a). **(G)** Most persistent H₁ loops identified in exposure and withdrawal networks. Serum metabolites are coloured by functional class: lipids (blue), amino acids and peptides (orange), carbohydrates and polyols (purple), acylcarnitines (red), organic acids and energy metabolites (green), and nucleotides and other small molecules (grey).

**Extended Data Figure 1.**
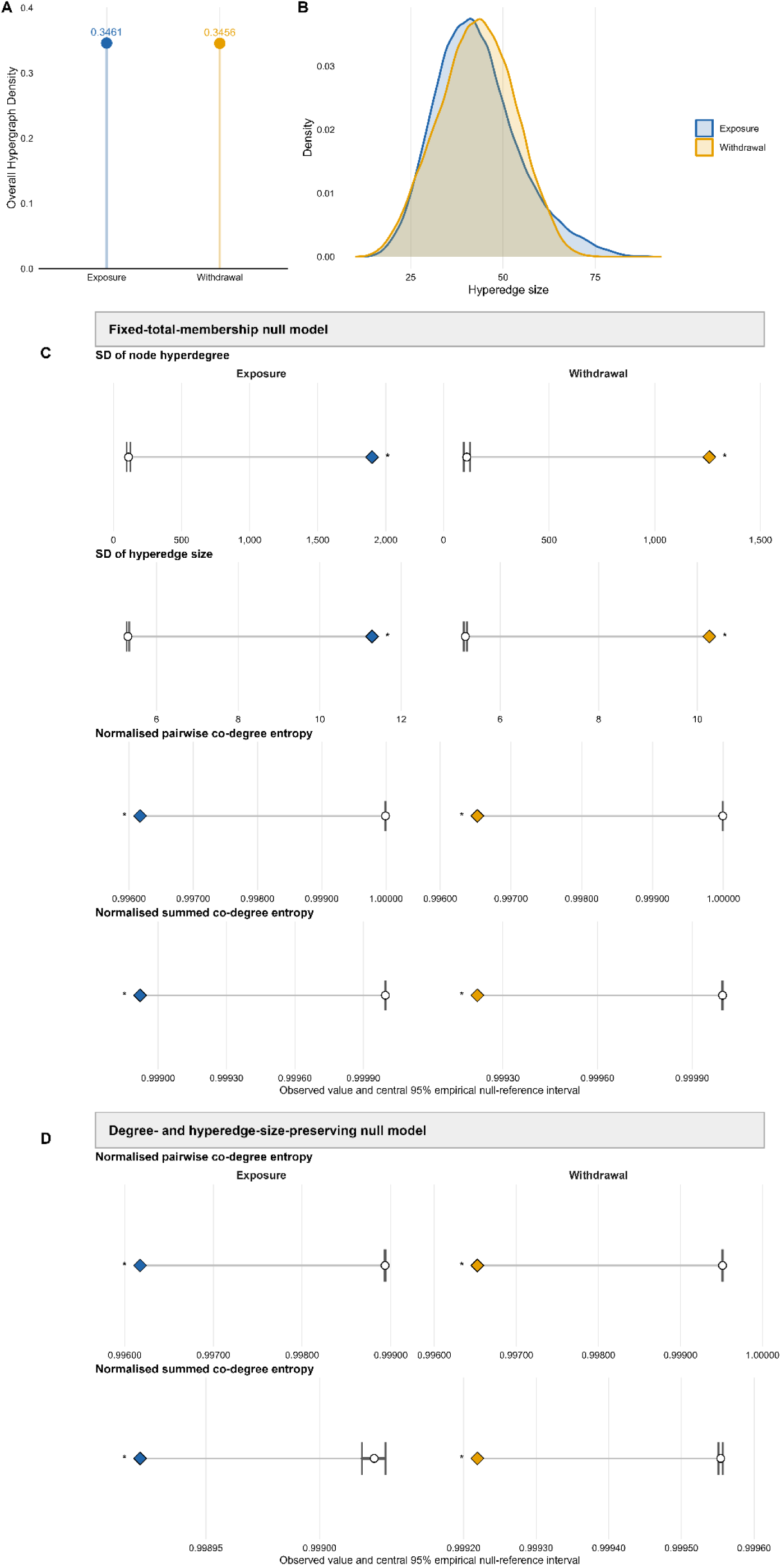
Global hypergraph architecture and null-model assessment. **(A)** Observed hypergraph density under GC exposure and withdrawal. (**B)** Hyperedge-size distributions, representing the number of serum metabolites associated with each adipose feature. (**C)** Structural metrics compared with a fixed-total-membership null model, which preserved total incidence and density. (**D)** Connectivity entropy compared with a degree-and hyperedge-size-preserving null model. (**C–D)** Diamonds show observed values, white circles show null medians, and error bars show central 95% empirical intervals from 1,000 randomisations. Pairwise co-degree entropy describes shared associations across metabolite pairs, whereas summed co-degree entropy describes shared connectivity across metabolites. Lower entropy indicates more concentrated connectivity. Blue denotes exposure and orange denotes withdrawal. Asterisks denote FDR-adjusted directional q-value.

Null-model analyses supported the non-random nature of this architecture. Under fixed-total-membership randomisation, exposure and withdrawal hypergraphs showed substantially greater node and hyperedge heterogeneity than expected by chance (Extended Data Fig. 1c). Under the more constraint degree- and hyperedge-size-preserving null model, degree-related metrics were preserved by design, but pairwise co-degree entropy and summed co-degree entropy remained significantly lower than expected in both states (Extended Data Fig. 1d). This indicates that shared-connectivity structure was non-random even after preserving marginal degree distributions. This provides evidence that the hypergraphs reflect genuine biological organisation rather than modelling artefacts.

#### GC exposure shifts metabolic networks towards higher-order coordination

Having established network-level differences in direct and shared connectivity, we next examined their relative contributions at the level of individual serum metabolites. As shared connectivity can increase partly due to greater node hyperdegree, the two measures were min max normalised separately within each GC state. A Δ shared-connectivity score was then calculated for each metabolite as normalised shared connectivity minus normalised node hyperdegree.

A marked GC-dependent shift in Δ shared-connectivity distributions was observed (Fig. 5c). During GC exposure, the distribution was shifted towards increased contribution of higher-order interactions relative to direct pairwise associations, reflecting shared connectivity and coordination across tissues. In contrast, GC withdrawal networks showed a distribution consistent with dominance of more independent pairwise relationships with adipose features and reduced overlap between metabolites. These findings show that GC exposure altered the balance between direct and shared connectivity at the level of individual circulating metabolites.

#### Topological data analysis reveals GC-dependent reorganisation of adipose-serum networks

Having shown that GC state altered both modular organisation and shared-connectivity structure within the adipose-serum hypergraphs, we next used topological data analysis (TDA) to assess global network organisation across scales (Fig. 5d). TDA identified clear differences between GC exposure and withdrawal networks, with persistence features across connected components (H0), loops (H1) and higher-dimensional voids (H2) showing distinct state-dependent patterns (Fig. 5e). The magnitude of these differences was quantified using Wasserstein distances, which were significantly greater between GC exposure and withdrawal networks than within-state comparisons (Fig. 5f, Extended Data Fig. 2a), indicating robust state-dependent topological separation. These findings indicate that GC exposure and withdrawal are associated with distinct and measurable changes in the underlying network structure, reflecting reorganization of multi-omic interactions between the serum and adipose tissue.

GC withdrawal was associated with increased fragmentation, reflected by higher persistence of connected components (H0) (Extended Data Fig. 2b). This pattern suggests the presence of more distinct and stable metabolite clusters, consistent with reduced integration across the network. In contrast, GC exposure showed increased persistence of higher-order loop structure (Extended Data Fig. 2b), suggesting greater coordination among groups of circulating metabolites linked to adipose multi-omic features. Persistent H1 loop composition provided biochemical support for the topological differences between GC states. The most persistent GC exposure loop contained a heterogeneous mixture of phospholipids, carbon metabolites, amino-acid-related metabolites and signalling molecules, whereas the most persistent withdrawal loop was enriched for free fatty acids and acylcarnitines (Fig. 5g). This pattern extended beyond individual loops, as across persistent H1 features, exposure-associated loops showed broader metabolite-class representation, while withdrawal-associated loops were more lipid-dominated (Extended Data Fig. 2c, Supplementary Table 1). These findings support a shift towards more diverse higher-order adipose-serum organisation during GC exposure.

Differences in H2 features, reflected in Wasserstein distance (Fig. 5f) and persistence summary statistics (Extended Data Fig. 2b), further supported topological separation between GC exposure and withdrawal networks.

Together, these analyses indicate that GC state reorganises adipose-serum relationships across multiple scales. Structural and shared-connectivity metrics showed that GC exposure was associated with greater heterogeneity and increased shared adipose-linked connectivity among circulating metabolites. TDA provided complementary global evidence of state-dependent topology, with withdrawal showing more persistent component separation and exposure showing more diverse higher-order loop structure. Viewed alongside the adipose pathway-level entropy findings, these results indicate that GC exposure reorganises metabolism across scales, from the structure of adipose metabolic pathways to the higher-order coordination linking adipose tissue and the circulation.

**Extended Data Figure 2:**
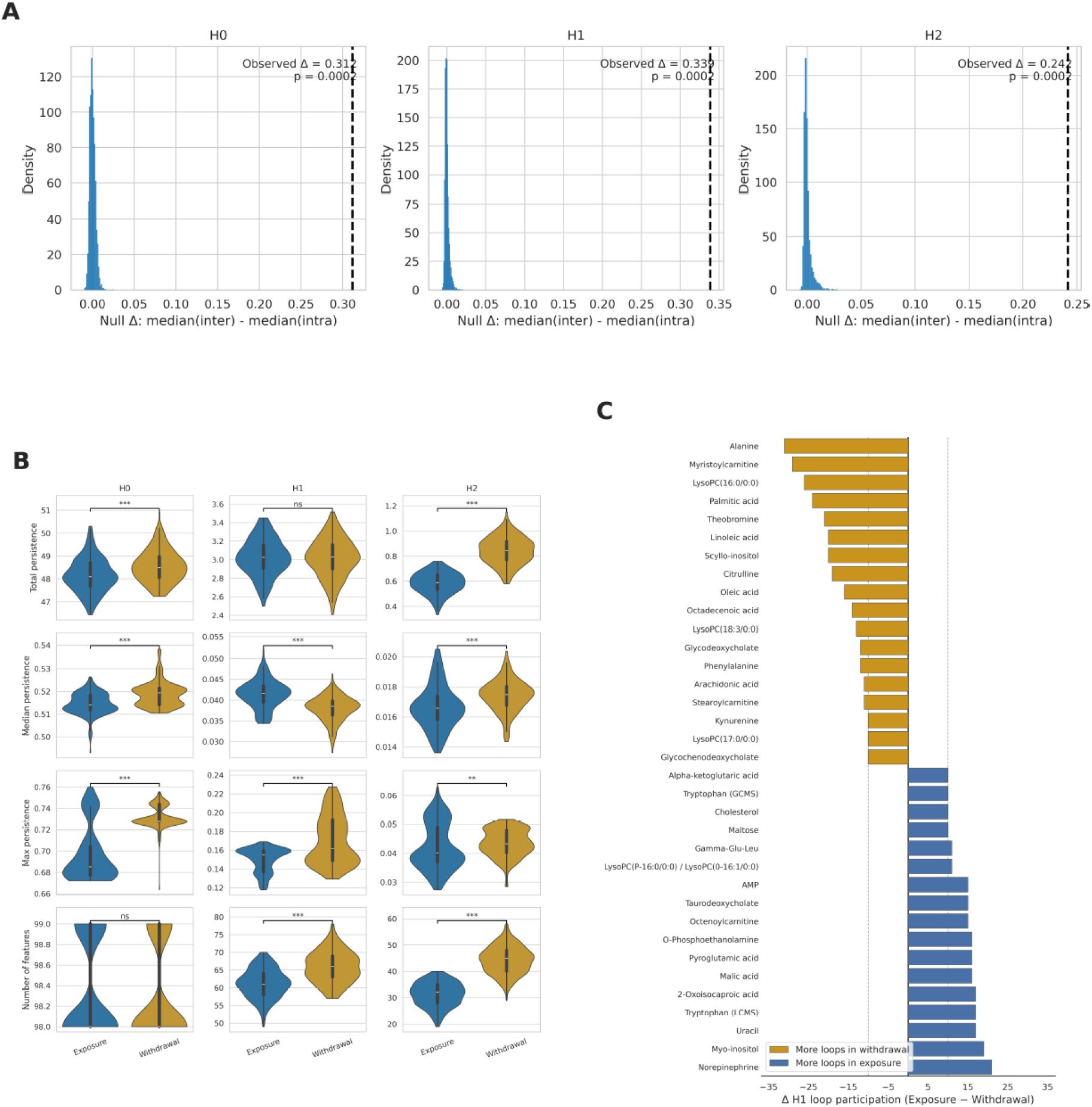
Permutation-based evidence and feature-level characterisation of GC-dependent topological reorganisation in adipose-serum networks. **(A)** Permutation-based assessment of topological separation between GC exposure and withdrawal networks. Null distributions (blue) show the expected difference in median Wasserstein distance between inter-condition and intra-condition persistence diagrams for homology dimensions H0-H2, generated by a label-based permutation test (5000 permutations). The dashed vertical line indicates the observed difference (Δ), demonstrating significantly greater topological dissimilarity between GC exposure and withdrawal than within-condition variability. **(B)** Topological differences between exposure and withdrawal persistence diagrams in homology dimensions H0, H1, and H2. Violin plots show the distribution of persistence diagram summary statistics across subsampled hypergraphs for exposure (blue) and withdrawal (yellow). Subsamples were generated by randomly selecting 80% of nodes (serum metabolites) without replacement (n = 200 subsamples per condition), and recomputing persistence diagrams for each subsample to assess robustness of topological features. Columns correspond to homology dimensions (H0–H2), and rows represent summary metrics including total persistence, median persistence, maximum persistence, and number of features. Statistical significance between conditions was assessed using permutation tests (two-sided, 5,000 permutations) on differences in group medians for each metric and dimension. **(C)** Persistent homology identifies metabolites differentially involved in network loops (H1) across GC states. Serum metabolites with a difference in the number of persistent loops (H1) between GC exposure and withdrawal networks of at least 10. Metabolites shown at the top in yellow appear in a greater number of loops during GC withdrawal, while those in blue are more frequently involved in loops during GC exposure.

### Identification of circulating markers reflecting adipose tissue GC action

#### Serum metabolites ratios outperform individual metabolites in discriminating GC state

Having established that GC exposure reorganises higher-order adipose-serum networks, we next assessed whether circulating metabolites could be used to distinguish GC exposure from withdrawal. Analysis of individual serum metabolites identified several candidates with strong predictive performance, including kynurenine, tryptophan and selected lipid species (Extended Data Table 3). However, given the coordinated, multi-pathway nature of GC-dependent metabolic reorganisation, we hypothesised that relational markers constructed from pairs of metabolites would better reflect GC state than univariate measures.

Metabolite ratios improved discriminatory performance, with multiple metabolite ratios achieving strong separation between GC exposure and withdrawal in this cohort (AUC 0.95-1, FDR ≤ 0.05; Fig. 6a, Supplementary Table 2). However, predictive performance alone does not establish whether a circulating marker reflects tissue-level GC effects. We therefore sought to distinguish markers that classify GC state from those that are mechanistically linked to GC-regulated processes within the adipose tissue.

**Extended Data Table 3:** Serum Metabolite univariate predictive markers between GC exposure and withdrawal at 7am (second intervention day). Abbreviations: LCMS, liquid chromatography mass spectrometry; GCMS, gas chromatography mass spectrometry.

|  | AUC | P-value | FDR |
| --- | --- | --- | --- |
| Kynurenine (GCMS) | 0.96 | 0.0004 | **0.016** |
| Tryptophan (LCMS) | 0.94 | 0.0005 | **0.016** |
| Kynurenine (LCMS) | 0.94 | 0.0002 | **0.016** |
| Tryptophan (GCMS) | 0.93 | 0.0005 | **0.016** |
| Docosahexaenoic Acid | 0.86 | 0.004 | 0.108 |
| Norepinephrine (noradrenaline) | 0.85 | 0.108 | 0.520 |
| Tyrosine (LCMS) | 0.83 | 0.015 | 0.300 |
| Phenylalanine | 0.82 | 0.057 | 0.499 |
| Hexadecanoic Acid | 0.81 | 0.049 | 0.482 |
| Linoleic Acid | 0.80 | 0.032 | 0.437 |
| Heptadecanoic Acid | 0.80 | 0.019 | 0.334 |
| Octadecadienoic Acid | 0.80 | 0.032 | 0.437 |
| Octadecenoic Acid | 0.80 | 0.078 | 0.520 |
| Tyrosine (GCMS) | 0.78 | 0.041 | 0.450 |
| Glutamine | 0.76 | 0.039 | 0.450 |

**Figure 6.**
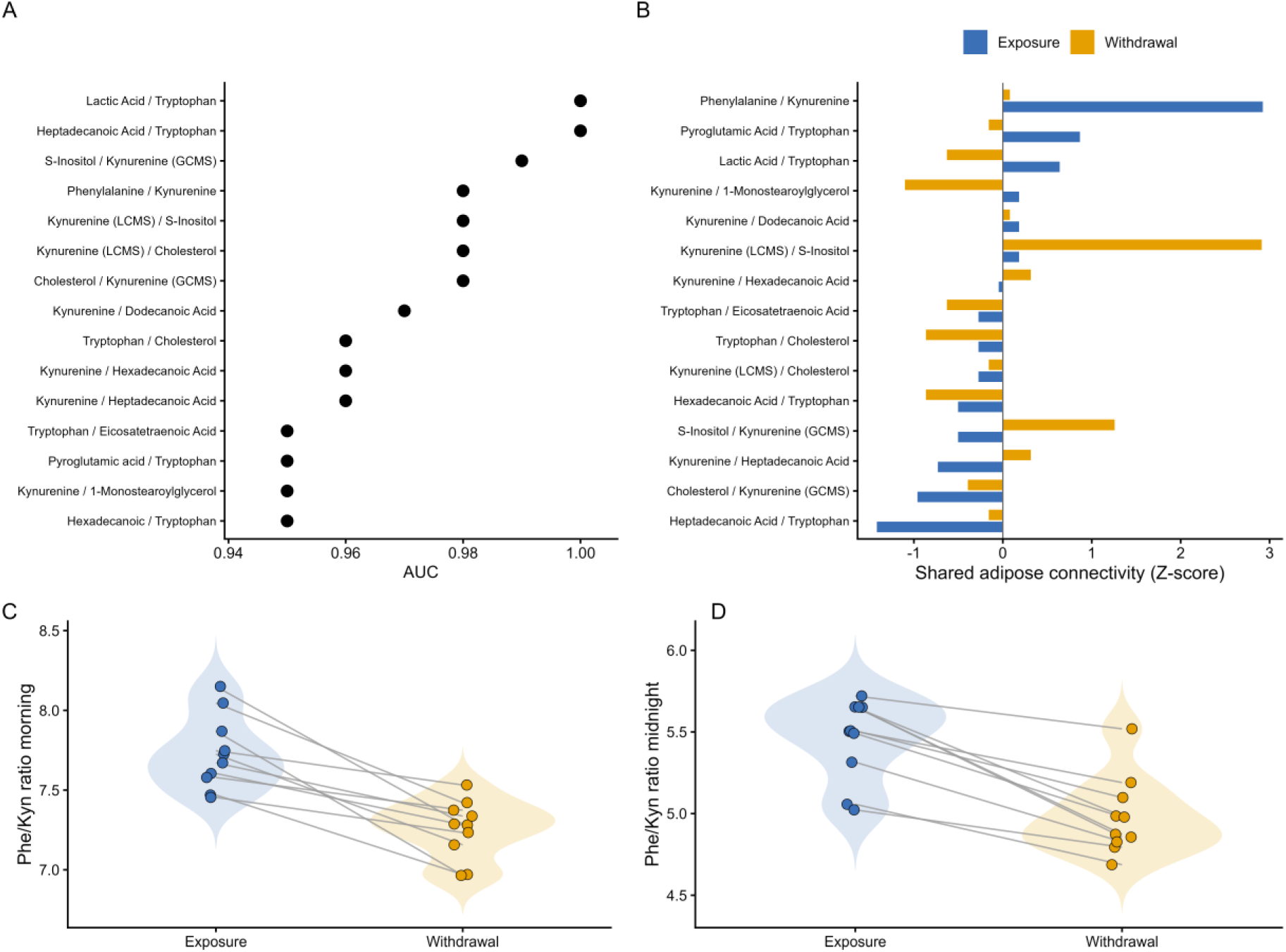
Integration of predictive performance and adipose network embedding identifies phenylalanine/kynurenine as a marker of GC state. **(A)** Predictive performance of the top 15 serum metabolite ratios for distinguishing GC exposure from GC withdrawal at 07:00. Metabolite ratios are ranked by area under the receiver operating characteristic curve (AUC). All displayed ratios remained significant following multiple-testing correction (FDR ≤ 0.05). **(B)** Enrichment of shared adipose tissue connectivity for the top 15 serum metabolite pairs. For each candidate metabolite pair, pairwise co-degree was calculated as the number of GC-regulated adipose transcripts and metabolites jointly associated with both serum metabolites. Pairwise co-degree values were standardised within each GC state and expressed as Z-scores for GC exposure (blue) and GC withdrawal (orange). Metabolite pairs are ranked by enrichment under GC exposure. **(C-D)** Serum phenylalanine/kynurenine ratio during GC exposure (blue) and GC withdrawal (orange). Paired measurements are shown for individual participants at **(C)** morning sampling (7am) corresponding to the physiological cortisol peak, and **(D)** midnight sampling, corresponding to the circadian trough.

#### Phenylalanine/kynurenine shows strongest adipose tissue connectivity in GC exposure networks

To prioritise circulating markers that reflect GC action in adipose tissue, we mapped the candidate circulating metabolite ratios onto the integrative adipose-serum hypergraph models. Within this hypergraph, we calculated the pairwise co-degree of the two metabolites forming each candidate ratio. Pairwise co-degree was defined as the number of GC-regulated adipose transcripts and metabolites jointly associated with both circulating metabolites, providing a measure of their shared embedding within GC-responsive adipose networks.

Among all candidate ratios, the phenylalanine/kynurenine ratio showed the strongest enrichment of shared associations with GC-regulated adipose tissue features during GC exposure (Z = 2.92; Fig. 6b; Supplementary Table 3). This indicates that these metabolites, although arising from distinct metabolic pathways, are jointly embedded within the GC-responsive adipose networks. Notably, the network position of this ratio exhibited the largest shift between GC exposure and withdrawal states, demonstrating dynamic reorganisation of its adipose connectivity in response to GC signalling. Hyperedges linking phenylalanine and kynurenine during GC exposure included well-established GC-response adipose genes, such as *TXNIP* and *KLF9*, supporting biological plausibility of this interaction.

In contrast, several other high-performing metabolite ratios showed weaker or non-specific adipose connectivity, suggesting that predictive performance alone is insufficient to identify biomarkers that reflect downstream tissue action. Integration of predictive performance with network context was therefore essential for prioritising biologically meaningful markers.

Consistent with this, individual metabolites contributing to the top-performing ratios, including phenylalanine, kynurenine and tryptophan, occupied distinct and GC-state-dependent positions within the shared-connectivity distributions (Extended Data Fig. 3). This reflects shifts in their higher-order connectivity to adipose tissue features. These findings suggest a network-level explanation for the improved performance of metabolite ratios: by combining metabolites whose interaction structure is reorganised by GC exposure, ratios capture changes in metabolic coordination rather than abundance alone.

Together, integration of predictive performance, hypergraph connectivity and interaction-order analyses identified the phenylalanine/kynurenine ratio as the circulating marker most strongly anchored to GC-regulated adipose tissue networks, providing a strong rationale for its evaluation across different GC exposure paradigms.

Within the BIOCORT crossover study, the phenylalanine/kynurenine ratio retained discriminatory power at the midnight sampling point, where serum cortisol levels remained higher during GC exposure than withdrawal (p<0.01 Wilcoxon signed-rank test), although this difference was attenuated compared to the morning peak. The ratio outperformed its constituent metabolites and other candidate metabolite ratios (AUC 0.90, FDR < 0.01, Supplementary Table 4). Importantly, the directionality of this effect was consistent with morning samples (Fig. 6c-d), indicating stability across circadian GC troughs.

**Extended Data Figure 3.**
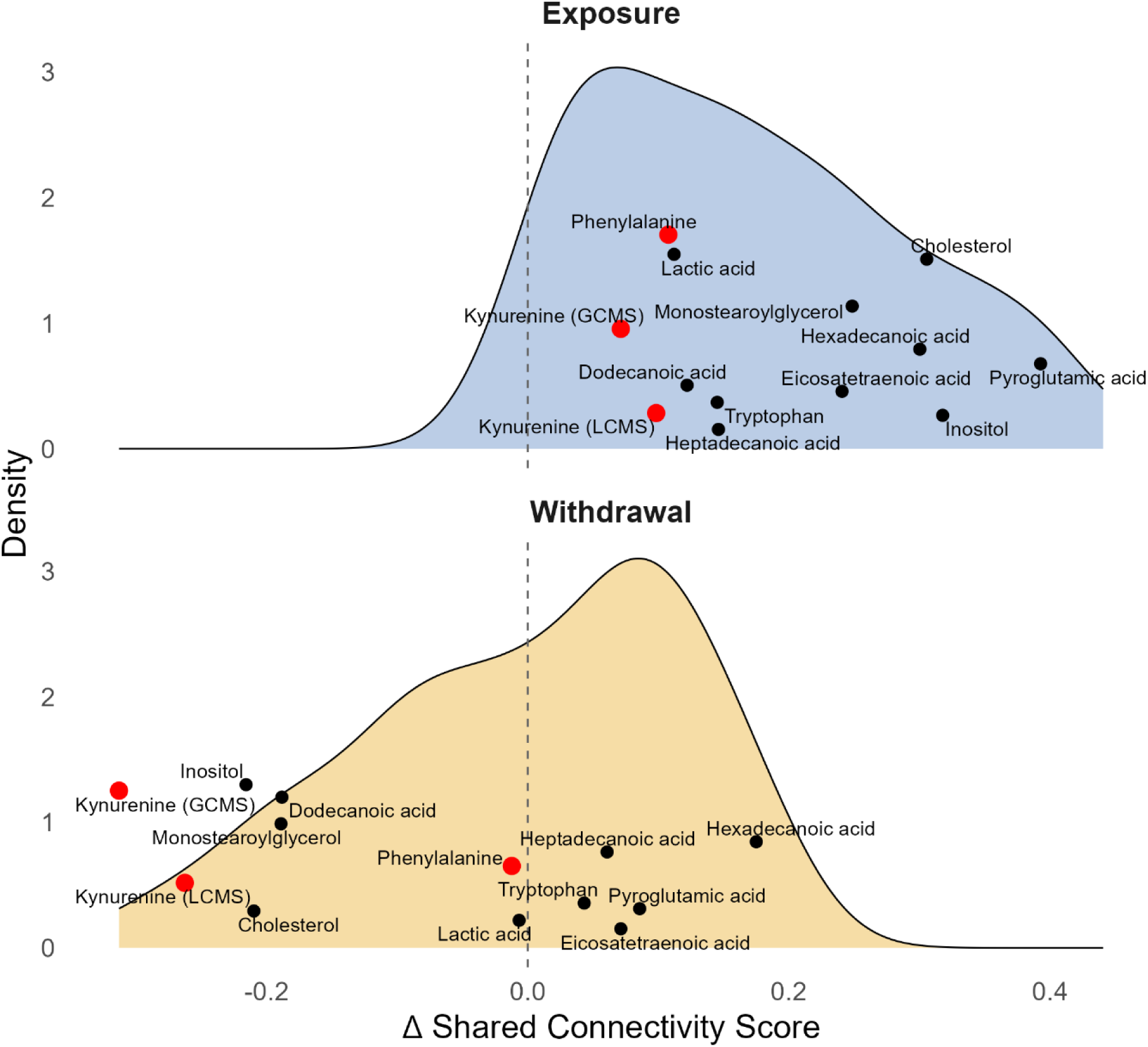
Position of candidate-marker metabolites within serum-adipose shared-connectivity distributions. Metabolites contributing to the candidate bivariate markers were mapped onto the condition-specific distributions of metabolite-level Δ shared connectivity during GC exposure (blue) and GC withdrawal (yellow). The Δ shared-connectivity score represents normalised summed co-degree minus normalised node hyperdegree, with positive values indicating disproportionately greater participation in shared adipose-linked associations relative to direct serum-adipose connectivity. Phenylalanine and kynurenine measurements contributing to the prioritised phenylalanine/kynurenine ratio are highlighted in red, while the remaining metabolites contributing to the evaluated candidate ratios are shown in black. The dashed vertical line indicates a Δ shared-connectivity score of zero.

#### Phenylalanine/kynurenine ratio replicates across independent GC exposure models

We next assessed the robustness of the phenylalanine/kynurenine ratio across independent experimental contexts with differing GC exposure profiles. In an independent dose-response study (DOSCORT), the ratio distinguished near-physiological from supra-physiological GC exposure following both acute (3h, p < 0.05) and prolonged (7 day, p < 0.01) treatment (Fig. 7). Notably, these effects were consistent in direction across the majority of the patients.

**Figure 7.**
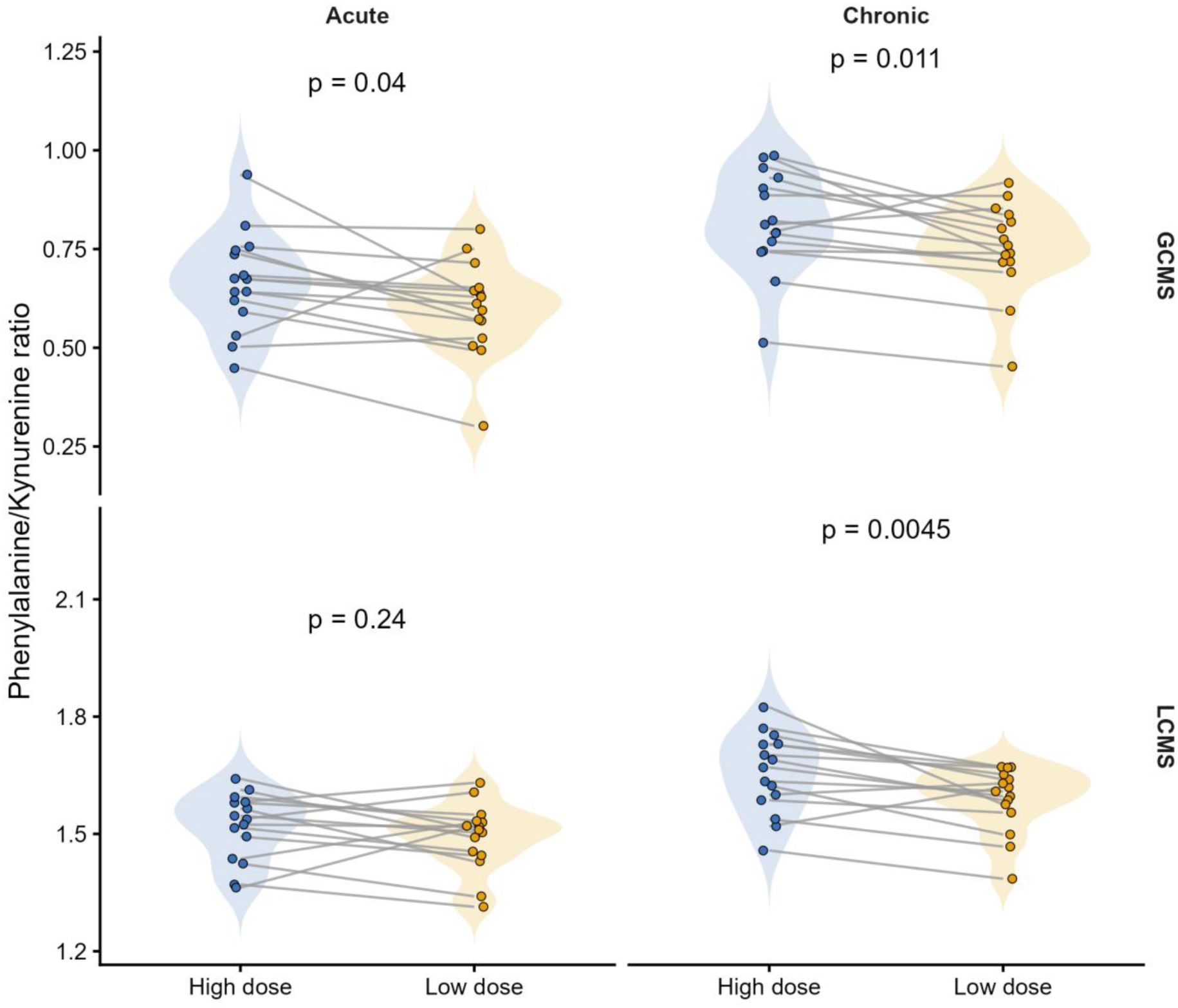
Ratio of phenylalanine/kynurenine as a marker of GC state in independent validation study. Phenylalanine/kynurenine ratio in serum after 3 hours (acute) and 7 days (chronic) of high dose (blue violin) vs low dose (orange violin) of betamethasone in DOSCORT study. The phenylalanine/kynurenine ratio in both GCMS (upper row) and LCMS (lower row) is shown. Doses were compared using a paired t-test, with P-values displayed.

Across both time points (3h and 7 days exposure), the phenylalanine/kynurenine ratio showed the strongest statistical association among all candidate markers (Supplementary Table 5). Although these associations did not remain significant following multiple-testing correction, the analysis was restricted to pre-specified candidate markers. This demonstrates sensitivity to both GC dose and duration of exposure.

Finally, in a separate study examining circadian GC exposure under different hydrocortisone dosing regimens^22,23^, the phenylalanine/kynurenine ratio exhibited significant 24-hour rhythmicity (Fig. 8a). While mean levels did not differ significantly between dosing strategies, the circadian pattern of the ratio was preserved across regimens. Kynurenine also exhibited rhythmic variation (Fig. 8a), whereas phenylalanine showed little evidence of rhythmicity, indicating that oscillations in the ratio were primarily driven by kynurenine.

**Figure 8.**
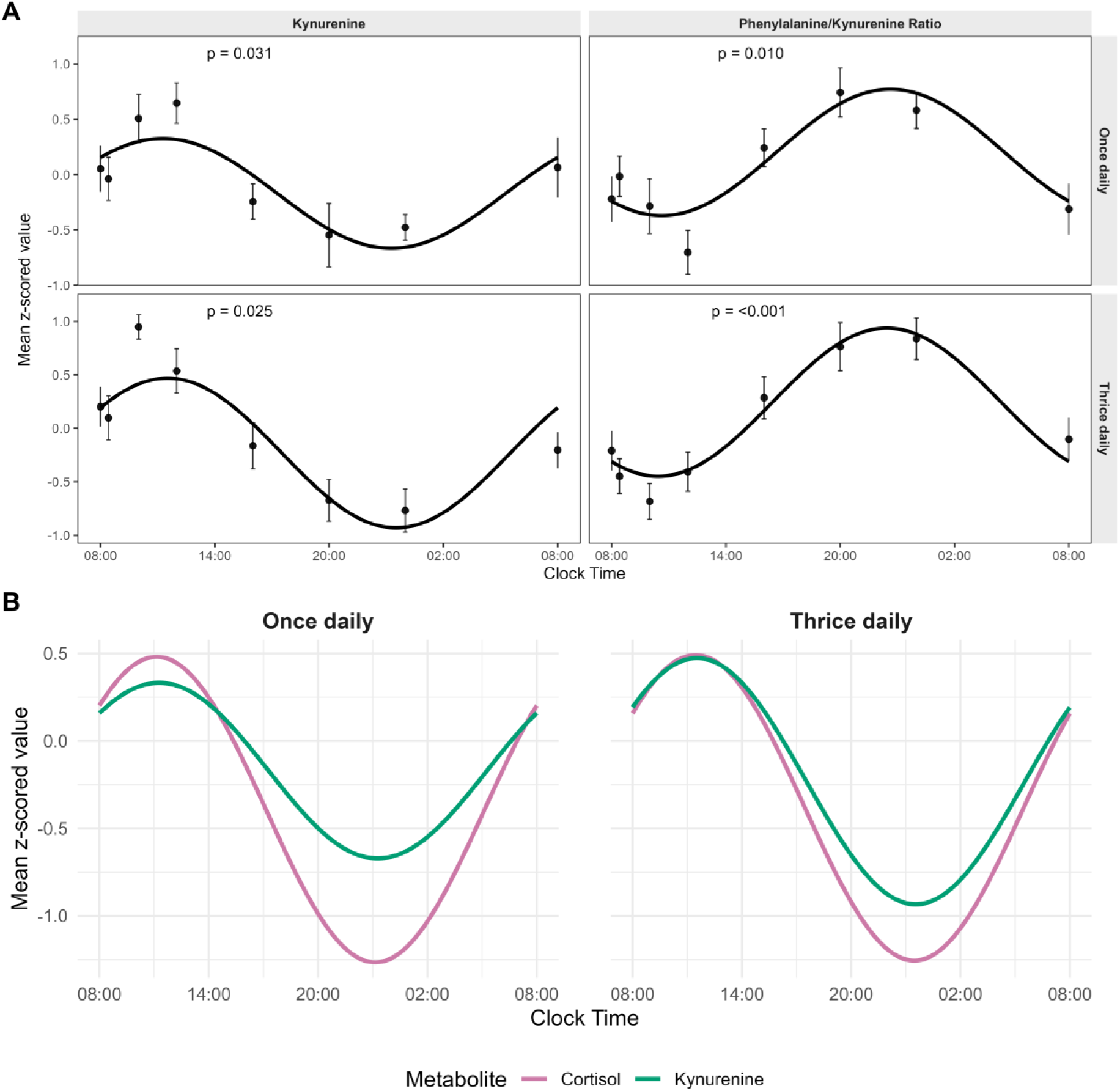
Circadian metabolite profiles under different dosing regimens. **(A)** Group-level cosinor fits for kynurenine and the phenylalanine/kynurenine ratio under once daily and thrice daily hydrocortisone dosing^22,23^. Regimens were matched for total daily oral dose but differed in temporal exposure profiles. Points represent mean z-scored metabolite values across the 24-hour sampling period, with error bars indicating standard error of the mean (SEM). Solid black lines show fitted group-level cosinor models. Top panels show once daily dual-release dosing, and bottom panels show thrice daily dosing. P-values indicate the statistical significance of the fitted cosinor model. **(B)** Overlay of cosinor fits for cortisol and kynurenine under once daily and thrice daily hydrocortisone dosing. Fitted curves are shown for mean z-scored cortisol (purple) and kynurenine (green) over 24 hours.

Cosinor modelling demonstrated broadly similar 24-hour phase patterns for cortisol and kynurenine under both dosing regimens (Fig. 8b). However, observed profiles indicated that this relationship was not strictly synchronous, with kynurenine peaking after cortisol, and overnight cortisol concentrations remaining low while kynurenine begins to rise (Extended Data Fig. 4). These findings support a temporal association between hydrocortisone exposure and kynurenine pathway activity, consistent with kynurenine reflecting integrated GC-sensitive metabolic regulation rather than instantaneous cortisol concentrations.

**Extended Data Figure 4.**
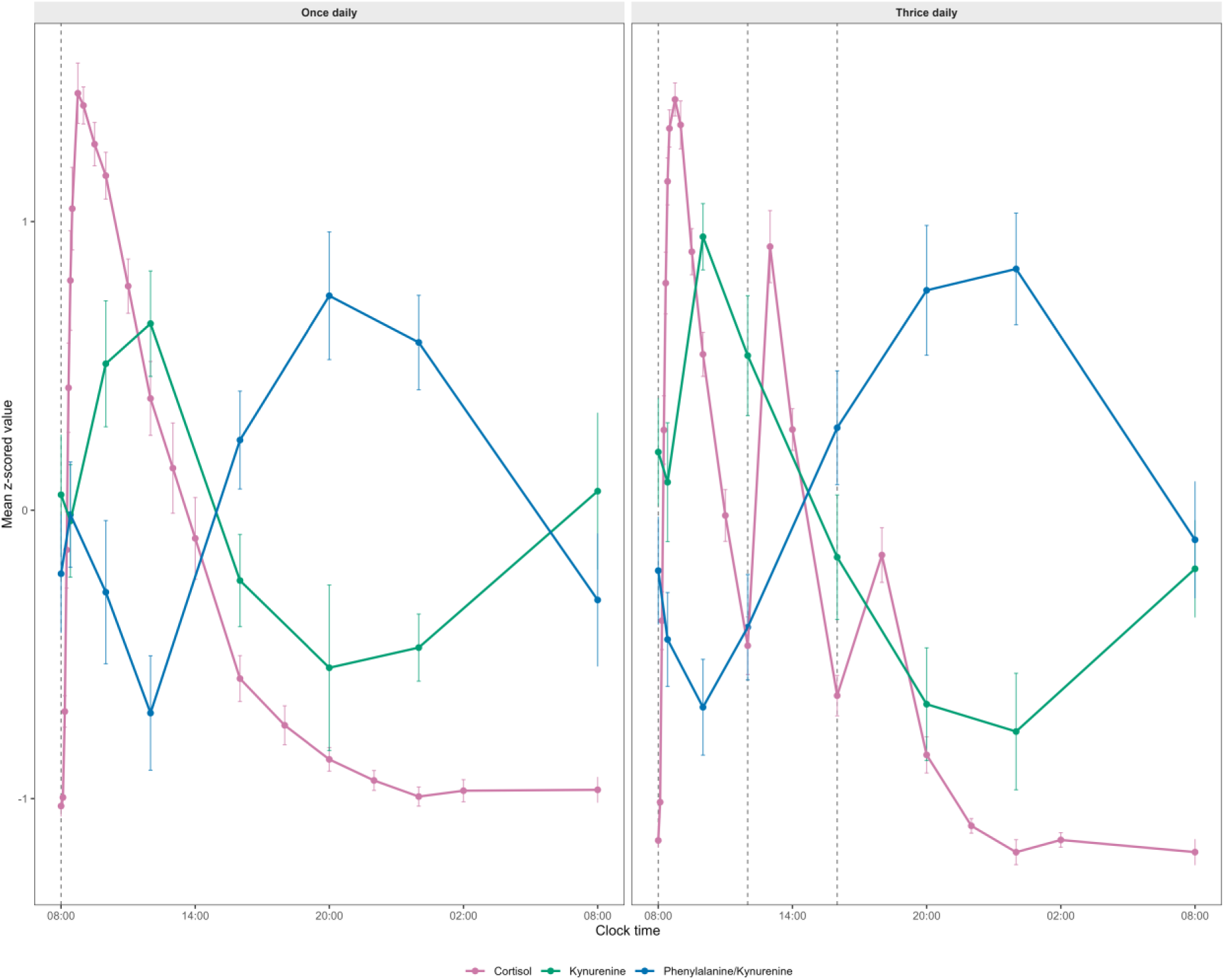
Observed 24-hour profiles of cortisol, kynurenine and phenylalanine/kynurenine ratio during hydrocortisone replacement. Mean (± SEM) within-subject z-scored values are shown for participants receiving once-daily (OD) and thrice-daily (TID) hydrocortisone replacement. Values are plotted at each sampling time point over 24 hours. Vertical dashed lines indicate hydrocortisone dosing times. Clock time is displayed relative to the first morning dose.

## DISCUSSION

Using a unique human model of controlled near-physiological GC exposure and withdrawal, we show that GCs not only remodel adipose tissue metabolism but also reorganise the higher-order networks linking adipose tissue to the circulation. These findings provide mechanistic insight into how tissue-specific GC action is mirrored in the circulation and identify the phenylalanine/kynurenine ratio as a circulating marker anchored to GC-regulated adipose networks.

Adipose tissue is a central target of the metabolic action of GCs, yet direct data on how physiological GC peaks and troughs regulate human adipose metabolism is limited. A key strength of this study is the use of in situ abdominal subcutaneous adipose tissue microdialysis to directly sample the interstitial metabolic environment in vivo, providing tissue-level resolution of local GC action. GC exposure was associated with coordinated changes in lipid and energy metabolism, including increased medium-chain acylcarnitines, consistent with enhanced fatty-acid flux coupled to incomplete mitochondrial β-oxidation in adipocytes. This aligns with evidence that GCs impair mitochondrial function and fatty-acid handling in hepatocytes and brown adipocytes^24,25^. Concurrent reductions in TCA-cycle intermediates and selective changes in phospholipid species suggest shifts in energy metabolism and membrane remodelling^26^. In parallel, changes in branched-chain amino acid, tryptophan-related and nucleoside metabolites indicate broader reprogramming of adipocyte metabolic state.

At the transcriptional level, GC exposure reorganised the higher-order connectivity within adipose metabolic pathways, rather than uniform transcriptional activation or repression. Entropy analysis revealed pathway-specific effects, with lipid and biosynthetic pathways showing reduced entropy during GC exposure, consistent with more focused and constrained transcriptional organisation, whereas carbohydrate and amino acid catabolic pathways showed the opposite pattern. This interpretation aligns with previous applications of entropy to biological networks. In signalling networks, higher entropy is associated with increased phenotypic plasticity and less constrained cellular states^27,28^, while reductions in local entropy can indicate more focused organisation within functionally specialised network regions^29^. These observations suggest that the circadian GC peaks and troughs impose distinct organisational states across different pathways of adipose metabolism.

At the systemic level, circulating metabolites are often interpreted as integrated readouts, strongly shaped by hepatic metabolism^30^, making tissue-specific effects difficult to disentangle. However, our paired adipose-serum analysis revealed substantial concordance under both GC states, alongside GC-dependent changes in how adipose metabolism is reflected in the circulation. Notably, kynurenine showed shifts in correlation direction and strength, indicating that its relationship with tissue metabolism is dynamically reconfigured by GC signalling, resulting in context-dependent coupling or decoupling between circulating and interstitial metabolite pools. Similar state-dependent changes in inter-tissue metabolite correlation structure have been observed in metabolic perturbation models^31^, emphasising the importance of inter-tissue relationships in interpreting circulating metabolomic profiles.

Comparison of hypergraphs with pairwise correlation-based networks showed only partial concordance in serum metabolite module assignments, indicating that higher-order modelling captures information beyond conventional pairwise associations^32,33^. Both hypergraph and topological analyses converged on the same biological signal: GC exposure promoted a more integrated and coordinated cross-tissue network, characterised by increased shared connectivity among circulating metabolites linked to common adipose features, whereas GC withdrawal was associated with network fragmentation and more independent adipose-serum associations. Together, these findings suggest that physiological GC exposure promotes coordinated cross-tissue metabolic regulation that is diminished during withdrawal.

Persistent H1 features suggested distinct modes of cross-tissue coordination, with withdrawal characterised by lipid-dominated loops and GC exposure involving broader contributions from phospholipids, amino acid derivatives and nucleotides. Because pathway entropy and topological analyses describe different aspects of metabolic organisation, these findings suggest that lipids may dominate cross-tissue coordination even when local adipose metabolic organisation shifts across pathways, consistent with their central role in inter-organ metabolic communication^34^. Overall, this shift from lipid-centred to multi-pathway coordination supports the interpretation that GC exposure promotes a more integrated metabolic state across tissues, whereas withdrawal favours a more compartmentalised organisation.

Within this framework, we sought circulating markers that reflect GC action within adipose tissue. Metabolite ratios outperformed individual metabolites in discriminating GC state, suggesting that relational markers better capture coordinated multi-pathway effects. Consistent with this, metabolite ratios can show stronger disease associations than their component metabolites^35^. Among all candidate ratios, the phenylalanine/kynurenine ratio demonstrated the strongest enrichment within GC-regulated adipose tissue networks, and the largest GC-state-dependent shift in network embedding. This indicates sensitivity to GC-dependent reorganisation of higher-order adipose-serum interactions. Several ratios with similarly high predictive performance showed limited adipose connectivity, highlighting the value of integrating discrimination and biological context for biomarker discovery.

Mechanistically, the phenylalanine/kynurenine ratio plausibly reflects GC-dependent regulation of amino acid metabolism across tissues. Kynurenine is generated through the tryptophan degradation pathway, regulated by hepatic tryptophan 2,3-dioxygenase (TDO) and extra-hepatic indoleamine 2,3-dioxygenase (IDO) in response to endocrine and inflammatory signals^36^. GCs have long been shown to induce hepatic TDO activity^37^, while adipose inflammation in obesity induces IDO^38^. Prior evidence that hydrocortisone alters tryptophan metabolism through the kynurenine pathway further supports the regulation of this axis by GCs^39^. Phenylalanine may reflect complementary changes in amino acid turnover, with adipose tissue contributing directly to systemic phenylalanine turnover^40^. Together, these observations suggest that phenylalanine/kynurenine reflects GC-dependent coordinated changes spanning hepatic amino acid metabolism, inflammatory signalling, and adipose metabolic state.

The robustness of this marker is supported by its consistency across experimental contexts. The phenylalanine/kynurenine ratio retained discriminatory power at the circadian trough and distinguished near-physiological from supra-physiological GC exposure in an independent dose-responsiveness study after both short-term and prolonged treatment. The ratio also showed significant circadian rhythmicity under different hydrocortisone replacement regimens. Notably, kynurenine exhibited significant rhythmicity, whereas phenylalanine showed little evidence of circadian variation, indicating that oscillations in the ratio were primarily driven by kynurenine. This is consistent with human evidence that kynurenine-pathway output varies across the day, with higher levels of kynurenine and related metabolites in the morning than at night^41^, and with animal and infection-model studies showing circadian regulation of kynurenine-pathway enzymes and IDO1-linked tryptophan metabolism^42,43^.

Although cosinor analyses suggested broad temporal alignment between cortisol and kynurenine rhythms, time-course profiles indicated that kynurenine peaks tended to follow post-dose cortisol rises, consistent with delayed regulation of the tryptophan-kynurenine pathway. This suggests that circulating kynurenine reflects integrated pathway activity over several hours rather than an immediate response to cortisol exposure. Consequently, circadian variation in the phenylalanine/kynurenine ratio may not fully predict its behaviour during sustained GC exposure, where longer-term changes in pathway flux and inter-tissue metabolism may alter the relationship between its component metabolites. Together, consistency across dose, duration and circadian phase indicates that the ratio reflects integrated GC action over time rather than transient metabolite fluctuations.

These findings have potential clinical implications for assessing GC exposure and tissue response. Current approaches rely largely on circulating cortisol concentrations, which do not capture tissue-specific sensitivity or downstream metabolic effects. A network-anchored biomarker such as phenylalanine/kynurenine may therefore offer a more physiologically relevant readout of GC action, addressing inter-individual variability in GC metabolism and tissue sensitivity, and supporting the development of personalised replacement therapy.

A major strength of this study is its direct investigation of GC action in humans. Although animal models have been instrumental in defining GC biology, species differences may limit translation of tissue-specific metabolic findings. The combination of controlled GC exposure, in vivo adipose microdialysis, transcriptomics and metabolomics in individuals lacking endogenous cortisol production provides a uniquely translatable framework for studying human GC action. However, several limitations should be considered. The sample size was modest (n=10), although the within-individual crossover design improved statistical efficiency and reduced inter-individual variation. The consistency of effects across participants, together with replication in independent cohorts with differing GC exposure paradigms, supports the robustness of the findings. As the BIOCORT study assessed short-term changes in GC exposure, longer-term studies will be required to determine how these network signatures relate to sustained metabolic outcomes. Moreover, although the paired design enables inference on adipose-serum coupling, it cannot fully resolve the organ-specific contributions to circulating metabolites. Validation in larger and more diverse cohorts will be required to confirm the predictive power of the phenylalanine/kynurenine ratio and define its mechanistic links to tissue-specific GC action.

In summary, our findings reveal the complex action of GCs in adipose tissue, where both peaks and troughs contribute to regulation of lipids, amino acids, and carbohydrate metabolism. Physiological GC exposure reorganises adipose metabolism and its coupling to the circulating metabolome through higher-order network structure. By integrating multi-omics data across tissues, we identify the phenylalanine/kynurenine ratio as a circulating marker linked to GC-regulated adipose networks. More broadly, these findings support a view of hormone action as the reorganisation of multi-tissue networks and provide a framework for developing biomarkers that reflect tissue-specific metabolic regulation.

## METHODS

### Clinical experimental study (<u>Bio</u>markers of <u>cort</u> isol-BIOCORT; *physiological circadian GC exposure vs withdrawal*)

#### Study design

This was a prospective, randomised, single-blind, two-period crossover clinical trial previously described in detail by Chantzichristos et al. (2020)^16^. Participants received either hydrocortisone or saline infusion in random order, with a washout period of at least two weeks between interventions. The study was approved by the Ethics Review Board of the University of Gothenburg (permit no. 374-13) and conducted in accordance with the Declaration of Helsinki. All patients received oral and written study information and signed informed consent before entering any part of the study. The trial was registered at ClinicalTrials.gov (NCT02152553).

#### Participants

Adults (20–60 years) with primary adrenal insufficiency (Addison’s disease) on stable hydrocortisone replacement therapy were recruited from the Centre for Adrenal Diseases, Sahlgrenska University Hospital, Gothenburg, Sweden. Inclusion and exclusion criteria, as well as recruitment procedures, have been described in detail by Chantzichristos et al. (2020)^16^. No formal power calculation was performed due to the exploratory nature of the study and the complexity of ‘omics’ analyses.

#### Study treatment

The hydrocortisone and saline infusion protocols have been described previously in detail^16^, based on earlier studies in healthy individuals^44^ and patients with adrenal insufficiency^45^. The hydrocortisone infusion was designed to mimic a physiological circadian cortisol rhythm, with a peak at 07:00 and a trough at midnight. During the withdrawal period, participants received an equal volume of 0.9% saline. Each intervention consisted of a 22-hour intravenous infusion of 2 litres for a 75 kg individual.

#### Interventions

Study procedures, including inpatient admission, standardised meals, and infusion protocols, have been described previously^16^. All morning samples were collected in a fasting state, following standardised meals the day before to reduce the impact of dietary variability. Caffeine intake (from coffee or tea) was recorded during the first intervention and replicated during the second intervention at identical times and quantities.

#### Sample Collection

Blood samples, adipose tissue biopsies, and microdialysis for interstitial metabolite profiling were obtained in that order starting at 07:00 to account for circadian variation. Blood samples at midnight (between first and second intervention day) were also collected and used for the replication study.

Abdominal subcutaneous adipose tissue samples were collected by aspiration. Local lidocaine was administered subcutaneously prior to sampling. Adipose biopsies were obtained from abdominal subcutaneous tissue and immediately snap-frozen, with more details provided in the Supplementary Methods.

Microdialysis of abdominal subcutaneous adipose tissue was performed to enable in vivo sampling of the interstitial fluid, providing a direct measurement of local metabolite dynamics and tissue-specific metabolic responses. Dialysate was collected over approximately 2 hours and stored at –80°C until analysis. Detailed methodological procedures are described in the Supplementary Methods.

#### Dialysate metabolic profiling by LCMS

Metabolic profiling of adipose tissue microdialysate was performed using liquid chromatography–mass spectrometry (LC–MS) at the Swedish Metabolomics Center (Umeå, Sweden). The LCMS workflow was optimised for detection of metabolites in adipose interstitial dialysate. Samples were analysed using an Agilent 1290 Infinity UHPLC system coupled to a 6546 Q-TOF mass spectrometer (Agilent Technologies, Waldbronn, Germany), operated in positive and negative electrospray ionisation modes. Metabolite identification was based on accurate mass, retention time and MS/MS fragmentation spectra, matched against in-house libraries generated from authentic standards analysed under identical conditions. Stable isotope-labelled internal standards were included during sample preparation, and samples were analysed in randomised order to minimise technical variation. Sample preparation, chromatographic conditions, and instrument parameters are described in Supplementary Methods.

#### Serum metabolomic profiling

Serum metabolomics was performed at the Swedish Metabolomics Center (Umeå, Sweden) using complementary gas chromatography–mass spectrometry (GC–MS) and liquid chromatography–mass spectrometry (LC–MS) platforms, as previously described^16^. Samples were analysed in randomised order and stable isotope-labelled internal standards were included throughout sample preparation and analysis to minimise technical variation.

GC–MS profiling primarily targeted central carbon metabolites, amino acids, organic acids and fatty acids. Raw GC–MS data were exported from ChromaTOF software and processed using established Swedish Metabolomics Center workflows, including baseline correction, chromatographic alignment and metabolite deconvolution. Compound identification was based on retention index and mass spectral matching against in-house and reference spectral libraries.

LC–MS profiling was performed using ultra-high-performance liquid chromatography coupled to high-resolution mass spectrometry operated in positive and negative ionisation modes. Metabolite identification was based on accurate mass, retention time and MS/MS spectral matching against authentic standards contained within in-house reference libraries. Quality-control procedures were applied throughout data acquisition and processing.

#### Adipose tissue transcriptomic profiling

Total RNA was extracted from abdominal subcutaneous adipose tissue biopsies collected during both GC exposure and GC withdrawal. RNA quantity and quality were assessed prior to transcriptomic analysis. Gene-expression profiling was performed using microarray-based transcriptomics at the Array and Analysis Facility, Science for Life Laboratory (Uppsala Biomedical Center, Uppsala, Sweden), as previously described^16^.

#### Data Analysis

All preprocessing, correlation-based integration, hypergraph construction, entropy analysis, biomarker overlap analysis, and figure assembly were performed in R. Persistent-homology analyses for the serum–adipose topological workflow were performed in Python using precomputed distance matrices, and H1 loop visualisation used pre-exported class-representative files from the Eirene workflow.

#### Metabolomics Identification

Raw LCMS and GCMS data were processed using the Agilent Masshunter Profinder version B.10.0.2 (Agilent Technologies Inc., Santa Clara, CA, USA). Feature extraction was performed using a targeted workflow (Batch Targeted Feature Extraction), in which a predefined list of metabolites commonly detected in plasma and serum was used.

Metabolite identification was based on matching accurate mass (MS), fragmentation spectra (MS/MS), and retention time against an in-house LCMS library generated from authentic standards analysed under identical chromatographic and mass spectrometry conditions.

#### Data Processing and Statistical Analysis of Metabolomics Data

Metabolomics data was analysed using Metaboboanalyst 6.0^46^. Duplicated features were removed, missing values were replaced by the limit of detection, and low-variance features were filtered. Preprocessing decisions were made on a data-set specific basis. Adipose microdialysate and DOSCORT metabolomics datasets were sum normalised due to variability in total metabolite signal between samples. BIOCORT serum metabolomics data, including the midnight validation samples, were analysed without sample-wise normalisation following quality-control assessment. DUOCORT metabolomics data were analysed using the previously established preprocessing framework for that dataset, including probabilistic quotient normalisation (PQN)^23^. All data was log transformed. Paired t-tests were used to assess differences between conditions, with statistical significance interpreted using both nominal P-values (P < 0.05), and false-discovery rate (FDR) adjustment was applied using the Benjamini-Hochberg method (FDR < 0.05). Volcano plots display log fold changes against -log10(P-value), with significance defined as P < 0.05 for visualisation.

#### Transcriptomic Data Processing and Statistical Analysis

Transcriptomic data from adipose tissue biopsies and PBMCs was analysed using the R packages *oligo*^47^ and *limma*^48^. Robust Multichip Average (RMA) was used for background subtraction, quantile normalisation and summarisation. Differentially expressed genes were found using linear modelling with empirical Bayes moderation. Genes with P-values < 0.05, and FDR < 0.05 were recorded.

#### Multivariate Analysis

Principal component analysis (PCA) was performed in R on adipose microdialysate metabolomics data after metabolite-wise mean-centering across samples. PCA was run using *prcomp* on the transposed data. The proportion of variance explained by each principal component (PC) was calculated from the PCA singular values, and the minimum number of components explaining at least 80% cumulative variance was retained, with a minimum of three components included. Associations between PC scores and selected clinical (e.g. age, sex, BMI, biochemical measures) and adipose transcriptomic variables (*CD68, ADIPOQ, SLC2A4*) were assessed using Pearson correlation for continuous variables and linear-model ANOVA for categorical variables. Effect sizes were reported as *r*^2^ or *n*^2^ respectively. Supervised multivariate classification was performed using partial least squares discriminant analysis (PLS-DA) implemented using the *mixOmics* package^49^.

#### Hypergraph Modelling

To characterise higher-order relationships within and between molecular datasets, we represented the data using hypergraphs, which extend conventional graphs by allowing one edge to connect more than two nodes simultaneously. Higher-order network approaches provide a framework for representing biological interactions that cannot be fully captured by pairwise relationships alone^33^. In a standard graph, an edge links only two nodes and therefore captures pairwise relationships. In contrast, a hypergraph can represent shared associations among groups of molecules, making them well suited to biological systems, in which multiple metabolites or transcripts may be associated with the same molecular feature^50^.

In this study, the definition of nodes and hyperedges depended on the analytical context. For pathway-level analysis of adipose transcriptomics, nodes represented genes within a given metabolic pathway, and hyperedges captured their shared associations with the wider transcriptome. For cross-tissue integration, nodes represented circulating metabolites, and hyperedges represented adipose tissue metabolites and transcripts. This approach allowed us to examine how groups of circulating metabolites collectively relate to adipose tissue features, capturing coordinated multi-omic structure across tissues.

Hypergraphs were constructed using correlation-based thresholding, and separately for each GC state to enable direct comparison of network organisation under GC exposure and withdrawal.

#### Pairwise versus Hypergraph Module Analysis

To assess whether hypergraph organisation could be reproduced by conventional pairwise representations, serum metabolite modules derived from hypergraphs were compared with modules obtained from three alternative network representations: a serum-metabolite correlation graph, a joint serum-adipose correlation graph, and a bipartite serum-adipose incidence representation. Modules were identified using hierarchical clustering and compared using the Adjusted Rand Index (ARI). Normalised Mutual Information (NMI), and Jaccard overlap were used as additional measures of module similarity.

#### Hypergraph Structural Analysis

Hypergraph structure was quantified from the binary incidence matrix. Node hyperdegree was calculated as the number of hyperedges connected to each serum metabolite (row sums of the incidence matrix). Hyperedge size was calculated as the number of serum metabolites associated with each adipose feature (column sums). A projected adjacency matrix was generated as:

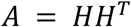

where *H* is the incidence matrix.

Before downstream calculations, diagonal entries were set to zero to remove self-counting of node hyperdegree. Each off-diagonal entry *A_ij_* represents the pairwise co-degree of serum metabolites *i* and *j*, defined as the number of adipose features jointly associated with both metabolites. For each serum metabolite, shared connectivity (*s_i_*) was calculated as the row sum of the off-diagonal matrix, equivalent to its summed co-degree with all other serum metabolites:

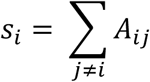

Shannon entropy was calculated both across the off-diagonal pairwise co-degree values and across the metabolite-level summed co-degree values. These measures are referred to as pairwise co-degree entropy and summed co-degree entropy, respectively, and were compared with corresponding null distributions.

#### Null Models

Statistical significance of both hypergraph module organisation and hypergraph structural properties was assessed using two complementary null models. A fixed-membership null model preserved the total number of node-hyperedge memberships while randomising their allocation within the incidence matrix. A second degree- and hyperedge-size preserving null model used Curveball-style randomisation to preserve node hyperdegree and hyperedge-size distributions while randomising higher-order structure. For each GC state, 1,000 null hypergraphs were generated. Observed module-similarity and structural metrics were compared with null distributions, and empirical P-values were adjusted using the Benjamini-Hochberg procedure.

#### Shared Connectivity Analysis

To quantify how circulating metabolites participate in shared associations with adipose tissue, we derived measures of connectivity within the hypergraph framework. Node hyperdegree reflects the number of adipose features linked to each metabolite and captures direct pairwise associations.

To assess higher-order coordination, we quantified shared connectivity as the summed co-degree of each serum metabolite with all other serum metabolites. This represents the extent to which metabolites participate in overlapping sets of adipose-linked associations, and provides a measure of coordinated network structure beyond individual pairwise relationships.

To compare GC states, both measures were min-max normalised within each condition, and their difference (Δ shared-connectivity score) was used to summarise the balance between higher-order and pairwise associations:

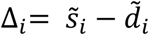

Where 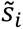 is shared connectivity, and 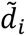 is node hyperdegree. Positive values indicate greater participation in shared, higher-order structures, whereas negative values indicate more independent, pairwise relationships. Differences in Δ shared-connectivity distributions between GC exposure and withdrawal were used to assess shifts in cross-tissue coordination.

#### Marker-Pair Enrichment Analysis

For biomarker-focused cross-tissue analyses, the adipose tissue feature set was restricted to GC-regulated metabolites and transcripts, therefore linking candidate circulating metabolites specifically to GC-responsive adipose processes.

For each pair of candidate serum metabolites, we quantified the extent to which they shared adipose-linked associations within the hypergraph. The pairwise co-degree between metabolites reflects the extent to which metabolites are jointly embedded within GC-regulated adipose networks.

Shared connectivity was computed separately for GC exposure and withdrawal and standardised within each condition using Z-scores. This enabled identification of metabolite pairs showing strong and condition-specific integration within adipose tissue features.

#### Pathway Entropy Analysis

To assess GC-dependent changes in the organisation of adipose transcriptomic networks, entropy was computed on pathway-specific hypergraphs constructed from gene expression data. Shannon entropy was calculated as:

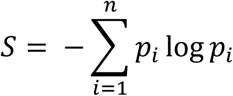

where:

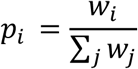

where *w_i_*represents co-degree values from the projected adjacency matrix.

Metabolic pathways were defined using Gene Ontology gene sets representing major branches of primary metabolism. Within each pathway, entropy was used to quantify how evenly higher-order shared associations were distributed across gene pairs. Lower entropy indicates that interactions are concentrated among a subset of genes, suggesting a more coordinated and constrained structure, whereas higher entropy reflects a more distributed organisation (Extended Data Fig. 5).

Pathway entropy was calculated separately for GC exposure and withdrawal and compared against size-matched random gene sets to control for transcriptome-wide effects. This allowed us to determine whether GC exposure induces pathway-specific reorganisation beyond global changes. Differences in entropy between GC states were assessed using Bayesian two-sample t-tests, because they enable comparison of the full entropy distributions rather than relying on summary statistics alone.

**Extended Data Figure 5.**
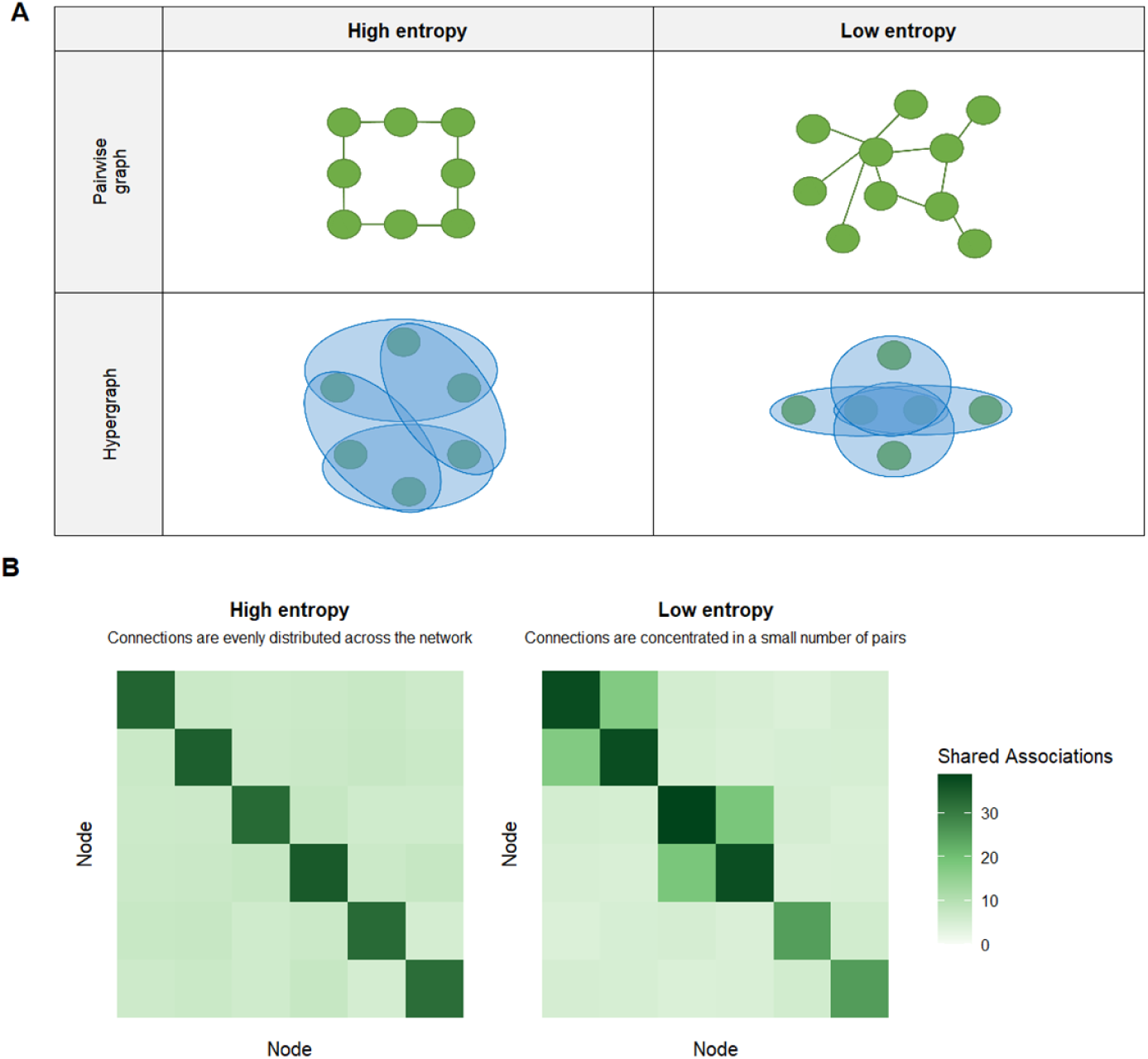
Conceptual illustration of entropy in pairwise and hypergraph representations. **(A)** Schematic illustration of entropy in pairwise graph and hypergraph representations. In pairwise graphs, nodes are connected by edges representing pairwise relationships. High entropy corresponds to an even distribution of connections, whereas low entropy reflects concentration of connections among a subset of nodes. In hypergraph representations, hyperedges connect groups of nodes, allowing higher-order relationships to be captured. Co-adjacency matrices derived from these hyperedges quantify how frequently pairs of nodes co-occur across interactions. High entropy reflects a uniform distribution of co-occurrence across many node pairs, whereas low entropy reflects concentration of interactions among a subset of node pairs. **(B)** Exemplar adjacency matrices illustrating high- and low-entropy configurations. Each matrix represents the number of shared associations between node pairs (e.g. the number of hyperedges in which two nodes co-occur). Both matrices contain the same total number of shared associations, but differ in how these associations are distributed across node pairs. High entropy is characterised by a relatively even distribution of shared associations across many node pairs, whereas low entropy reflects concentration of shared associations within a limited subset of node pairs. Colour scale indicates the magnitude of shared associations.

#### Topological Data Analysis

To characterise global network structure, we applied topological data analysis (TDA) to the serum-adipose hypergraphs. This approach identifies patterns of connectivity, loops, and higher-dimensional features, and has previously been used to identify biologically meaningful structure in complex molecular networks^51^.

Pairwise distances between metabolites in the hypergraph were derived from similarity in their adipose-linked association profiles, using the Jaccard index. This distance matrix was used to construct persistence diagrams describing topological features across increasing scales, including connected components (H0), loops (H1), and voids (H2).

Differences between GC exposure and withdrawal were quantified using Wasserstein distances between persistence diagrams, providing a measure of global topological similarity. To ensure robustness, analyses were repeated across sub-sampled networks, and statistical significance was assessed using a label-based permutation test. For persistence-summary analyses, each pseudo-replicate diagram was additionally summarised by the number of finite features, total persistence, median persistence, and maximum persistence in each homology dimension.

To interpret biochemical structure underlying higher-order network features, we analysed metabolite participation in persistent H1 loop structures identified through TDA. Metabolites contributing to these loops were identified and compared between GC exposure and withdrawal. Differences in loop participation were used to highlight metabolites and functional classes that contribute to higher-order coordination under each GC state.

#### Predictive Biomarker Analysis

In the discovery cohort, receiver operating characteristic (ROC) curve analysis within MetaboAnalyst was used to quantify the discriminatory performance of individual serum metabolites and metabolite ratios for distinguishing GC exposure from GC withdrawal. Area under the curve (AUCs), P-values and FDR-adjusted (Benjamini-Hochberg) P-values are reported. An AUC of 1 indicates complete discrimination between exposure and withdrawal groups. These discovery rankings were then integrated with the hypergraph-based overlap analyses described above to prioritise candidate circulating markers showing both discriminatory performance and embedding within GC-regulated adipose networks.

#### Cosinor Analysis

To evaluate circadian dynamics of candidate metabolite markers, we applied cosinor modelling to assess 24-hour rhythmicity. Metabolite and cortisol values in the DUOCORT study were standardised within individuals prior to fitting group-level models. This approach enabled quantification of rhythmic patterns and comparison of circadian dynamics across different GC exposure regimens.

#### Replication studies

To validate circulating markers identified in the BIOCORT discovery analysis, these were evaluated across complementary settings capturing variation in GC exposure, including circadian phase (BIOCORT midnight), dose-dependent effects (DOSCORT), and physiological circadian replacement (DUOCORT). Analyses were restricted to metabolites and metabolite ratios available in each dataset. Where equivalent metabolites were measured across both LCMS and GCMS platforms, platform-specific results were retained and analysed separately.

#### BIOCORT (midnight time point)

Candidate metabolite markers identified at 07:00 were re-evaluated at the midnight sampling time point within the BIOCORT study. Discriminatory performance was assessed using the same receiver operating characteristic (ROC) and area under the curve (AUC) framework applied in the discovery analysis, including both metabolite ratios and individual metabolites.

#### DOSCORT (dose-response study)

The DOSCORT study was a randomised, two-period, double-blind cross-over trial comparing low (0.5 mg) and high (1.25 mg) doses of betamethasone administered once daily for seven days, with a 2–5 week washout period between interventions. The study included 15 patients with Addison’s disease. Serum samples for metabolomic analysis were collected in the fasting state at baseline (prior to dosing), 3 hours after the first dose (acute exposure), and following 7 days of treatment (chronic exposure). Samples were stored at −80°C and analysed in a single batch. To assess dose-dependent effects of GC exposure, candidate metabolite markers were compared between low- and high-dose conditions at both the acute (3-hour) and chronic (7-day) time points. Paired t-tests were performed within each time point, and P-values were adjusted for multiple testing using the Benjamini–Hochberg method.

#### DUOCORT (circadian exposure study)

The DUOCORT study was a randomised, controlled, two-period cross-over trial comparing once-daily dual-release hydrocortisone (OD) with conventional hydrocortisone administered three times daily (TID)^22^, representing more and less physiological circadian GC exposure profiles, respectively. The study included 18 patients with Addison’s disease on stable GC replacement therapy.

Participants underwent 24-hour inpatient sampling at the end of each treatment period, with serum collected at eight time points under standardised conditions.

To evaluate circadian dynamics of candidate metabolite markers, metabolite concentrations were analysed using cosinor modelling. Values were z-score standardised within individuals prior to fitting group-level 24-hour models, enabling assessment of rhythmicity across dosing regimens.

## DATA AVAILABILITY

Data underlying this study have been deposited in public repositories. BIOCORT serum metabolomics datasets (LCMS and GCMS) are available through Mendeley Data (DOI: 10.17632/7hc49hzzhc.2). Adipose tissue transcriptomic data are available through the Gene Expression Omnibus (GEO) under accession GSE148642. The DUOCORT metabolomics dataset used in this study has been deposited in the MetaboLights repository under study identifier MTBLS13147. The adipose microdialysis metabolomics dataset generated during the current study are being deposited in MetaboLights, and accession numbers will be provided upon completion of the submission process. All code used for data processing, statistical analysis, hypergraph construction, topological data analysis and figure generation is being prepared for public release through GitHub. Repository details will be provided upon completion.

## ETHICS

The biomarker discovery study was registered at ClinicalTrials.gov under the identifier NCT02152553. It received ethical approval from the Ethics Review Board of the University of Gothenburg, Sweden (permit no. 374-13, August 8, 2013). The replication DOSCORT study was registered under the identifiers EudraCT: 2016-004078-16; ClinicalTrials.gov ID: NCT03210545. It received ethical approval from the Ethics Review Board of the University of Gothenburg, Sweden (permit no. 584-16) and was approved by the Swedish Medical Product Agency (MPA). The replication DUOCORT study was registered at EudraCT under the identifier 2006-0007084-83 and at ClinicalTrials.gov under the identifier NCT00915343. It received ethical approval from the Ethics Review Board of the University of Gothenburg, Sweden (permit no. 104-07 and T295-16). All studies were conducted in accordance with the Declaration of Helsinki. Written informed consent was obtained from all participants prior to their enrolment in the studies.

## ACKNOWLEDGEMENTS

We express our gratitude to all the three studies’ volunteers for their participation. We also extend our thanks to the following contributors to the project: the In-patient and Out-patient Clinics and the Endocrine Research Unit at the Department of Endocrinology-Diabetes-Metabolism, Sahlgrenska University Hospital, Gothenburg, Sweden,; research nurses Lena Strindberg and nurses Olof Ehn, Ingrid Broms, Frida Gillberg, Ann-Charlotte Olofsson, Kristina Cid Käll and Rebecka Starke; the Array and Analysis Facility, Science for Life Laboratory at Uppsala Biomedical Center (BMC), Uppsala, Sweden; Hans Stenlund at the Swedish Metabolomics Center, Umeå, Sweden; Ruth Andrew and Natalie Homer at the Mass Spectrometry Core Laboratory, Clinical Research Facility, University of Edinburgh, Edinburgh, UK; Professor Martin Schmelz, Dept of Anesthesiology and Intensive Care Medicine Mannheim, University of Heidelberg, Heidelberg, Germany for manufacturing of the microdialysis catheters.

## REFERENCES

1 Timmermans, S., Souffriau, J. & Libert, C. A General Introduction to Glucocorticoid Biology. Front Immunol 10, 1545 (2019). 10.3389/fimmu.2019.01545

2 Johannsson, G. et al. Adrenal insufficiency: review of clinical outcomes with current glucocorticoid replacement therapy. Clinical Endocrinology 82, 2–11 (2015). 10.1111/cen.12603

3 Dineen, R., Stewart, P. M. & Sherlock, M. Factors impacting on the action of glucocorticoids in patients receiving glucocorticoid therapy. Clinical Endocrinology 90, 3–14 (2019). 10.1111/cen.13837

4 Vegiopoulos, A. & Herzig, S. Glucocorticoids, metabolism and metabolic diseases. Mol Cell Endocrinol 275, 43–61 (2007). 10.1016/j.mce.2007.05.015

5 Bergthorsdottir, R. et al. Increased risk of hospitalization, intensive care and death due to COVID-19 in patients with adrenal insufficiency: A Swedish nationwide study. Journal of Internal Medicine 295, 322–330 (2024). 10.1111/joim.13731

6 Dalakas, K. et al. Increased mortality in primary adrenal insufficiency: a systematic review and meta-analysis. European Journal of Endocrinology 193, S53–S60 (2025). 10.1093/ejendo/lvaf186

7 Peckett, A. J., Wright, D. C. & Riddell, M. C. The effects of glucocorticoids on adipose tissue lipid metabolism. Metabolism 60, 1500–1510 (2011). 10.1016/j.metabol.2011.06.012

8 Lee, M. J., Pramyothin, P., Karastergiou, K. & Fried, S. K. Deconstructing the roles of glucocorticoids in adipose tissue biology and the development of central obesity. Biochim Biophys Acta 1842, 473–481 (2014). 10.1016/j.bbadis.2013.05.029

9 Lee, R. A., Harris, C. A. & Wang, J. C. Glucocorticoid Receptor and Adipocyte Biology. Nucl Receptor Res 5 (2018). 10.32527/2018/101373

10 Akalestou, E., Genser, L. & Rutter, G. A. Glucocorticoid Metabolism in Obesity and Following Weight Loss. Front Endocrinol (Lausanne) 11, 59 (2020). 10.3389/fendo.2020.00059

11 Oster, H. et al. The Functional and Clinical Significance of the 24-Hour Rhythm of Circulating Glucocorticoids. Endocr Rev 38, 3–45 (2017). 10.1210/er.2015-1080

12 Gooley, J. J. Circadian regulation of lipid metabolism. Proc Nutr Soc 75, 440–450 (2016). 10.1017/S0029665116000288

13 Zhang, Z., Xin, H. & Li, M. D. Circadian Rhythm of Lipid Metabolism in Health and Disease. Small Methods 4 (2020). 10.1002/smtd.201900601

14 Tholen, S. et al. Flattening of circadian glucocorticoid oscillations drives acute hyperinsulinemia and adipocyte hypertrophy. Cell Rep 39, 111018 (2022). 10.1016/j.celrep.2022.111018

15 Cain, D. W. & Cidlowski, J. A. Immune regulation by glucocorticoids. Nat Rev Immunol 17, 233–247 (2017). 10.1038/nri.2017.1

16 Chantzichristos, D. et al. Identification of human glucocorticoid response markers using integrated multi-omic analysis from a randomized crossover trial. Elife 10 (2021). 10.7554/eLife.62236

17 Kleeman, S. O. et al. Cystatin C is glucocorticoid responsive, directs recruitment of Trem2+ macrophages, and predicts failure of cancer immunotherapy. Cell Genom 3, 100347 (2023). 10.1016/j.xgen.2023.100347

18 Kern, P. A., Di Gregorio, G. B., Lu, T., Rassouli, N. & Ranganathan, G. Adiponectin expression from human adipose tissue: relation to obesity, insulin resistance, and tumor necrosis factor-alpha expression. Diabetes 52, 1779–1785 (2003). 10.2337/diabetes.52.7.1779

19 Kouidhi, S. et al. Human subcutaneous adipose tissue Glut 4 mRNA expression in obesity and type 2 diabetes. Acta Diabetol 50, 227–232 (2013). 10.1007/s00592-011-0295-8

20 Di Gregorio, G. B. et al. Expression of CD68 and Macrophage Chemoattractant Protein-1 Genes in Human Adipose and Muscle Tissues: Association With Cytokine Expression, Insulin Resistance, and Reduction by Pioglitazone. Diabetes 54, 2305–2313 (2005). 10.2337/diabetes.54.8.2305

21 Stifel, U. et al. Glucocorticoids coordinate macrophage metabolism through the regulation of the tricarboxylic acid cycle. Mol Metab 57, 101424 (2022). 10.1016/j.molmet.2021.101424

22 Johannsson, G. et al. Improved cortisol exposure-time profile and outcome in patients with adrenal insufficiency: a prospective randomized trial of a novel hydrocortisone dual-release formulation. J Clin Endocrinol Metab 97, 473–481 (2012). 10.1210/jc.2011-1926

23 McQueen, J. et al. Different metabolic pathways associated with total cortisol exposure and the cortisol time profile: a randomized crossover trial. Scientific Reports 16, 6510 (2026). 10.1038/s41598-026-37816-0

24 Lettéron, P. et al. Glucocorticoids inhibit mitochondrial matrix acyl-CoA dehydrogenases and fatty acid beta-oxidation. Am J Physiol 272, G1141–1150 (1997). 10.1152/ajpgi.1997.272.5.G1141

25 Tang, X. et al. Dexamethasone-induced whitening of rabbit brown adipose tissue: leptin resistance and mitochondrial dysfunction. BMC Genomics 26, 326 (2025). 10.1186/s12864-025-11502-3

26 Cho, Y. K. et al. Lipid remodeling of adipose tissue in metabolic health and disease. Exp Mol Med 55, 1955–1973 (2023). 10.1038/s12276-023-01071-4

27 Teschendorff, A. E. & Severini, S. Increased entropy of signal transduction in the cancer metastasis phenotype. BMC Syst Biol 4, 104 (2010). 10.1186/1752-0509-4-104

28 Banerji, C. R. et al. Cellular network entropy as the energy potential in Waddington’s differentiation landscape. Sci Rep 3, 3039 (2013). 10.1038/srep03039

29 West, J., Bianconi, G., Severini, S. & Teschendorff, A. E. Differential network entropy reveals cancer system hallmarks. Sci Rep 2, 802 (2012). 10.1038/srep00802

30 Wu, Z. E. et al. Dissecting the relationship between plasma and tissue metabolome in a cohort of women with obesity: Analysis of subcutaneous and visceral adipose, muscle, and liver. FASEB J 36, e22371 (2022). 10.1096/fj.202101812R

31 Sato, S. et al. Atlas of exercise metabolism reveals time-dependent signatures of metabolic homeostasis. Cell Metab 34, 329–345 e328 (2022). 10.1016/j.cmet.2021.12.016

32 Gaudelet, T., Malod-Dognin, N. & Przulj, N. Higher-order molecular organization as a source of biological function. Bioinformatics 34, i944–i953 (2018). 10.1093/bioinformatics/bty570

33 Battiston, F. et al. The physics of higher-order interactions in complex systems. Nature Physics 17, 1093–1098 (2021). 10.1038/s41567-021-01371-4

34 Liu, S., Alexander, R. K. & Lee, C. H. Lipid metabolites as metabolic messengers in inter-organ communication. Trends Endocrinol Metab 25, 356–363 (2014). 10.1016/j.tem.2014.05.002

35 Molnos, S. et al. Metabolite ratios as potential biomarkers for type 2 diabetes: a DIRECT study. Diabetologia 61, 117–129 (2018). 10.1007/s00125-017-4436-7

36 Xue, C. et al. Tryptophan metabolism in health and disease. Cell Metab 35, 1304–1326 (2023). 10.1016/j.cmet.2023.06.004

37 Feigelson, P. & Greengard, O. Immunologic and enzymatic studies on rat liver tryptophan pyrrolase during substrate and cortisone induction. Ann N Y Acad Sci 103, 1075–1082 (1963). 10.1111/j.1749-6632.1963.tb53759.x

38 Favennec, M. et al. The kynurenine pathway is activated in human obesity and shifted toward kynurenine monooxygenase activation. Obesity (Silver Spring) 23, 2066–2074 (2015). 10.1002/oby.21199

39 Sorgdrager, F. J. H., Werumeus Buning, J., Bos, E. H., Van Beek, A. P. & Kema, I. P. Hydrocortisone Affects Fatigue and Physical Functioning Through Metabolism of Tryptophan: A Randomized Controlled Trial. J Clin Endocrinol Metab 103, 3411–3419 (2018). 10.1210/jc.2018-00582

40 Coppack, S. W., Persson, M. & Miles, J. M. Phenylalanine kinetics in human adipose tissue. J Clin Invest 98, 692–697 (1996). 10.1172/jci118840

41 Rapoport, M. I. & Beisel, W. R. Circadian periodicity of tryptophan metabolism. J Clin Invest 47, 934–939 (1968). 10.1172/jci105785

42 Moravcová, S. et al. Circadian control of kynurenine pathway enzymes in the rat pineal gland, liver, and heart and tissue- and enzyme-specific responses to lipopolysaccharide. Arch Biochem Biophys 722, 109213 (2022). 10.1016/j.abb.2022.109213

43 Stincardini, C. et al. The circadian control of tryptophan metabolism regulates the host response to pulmonary fungal infections. PNAS Nexus 2, pgad036 (2023). 10.1093/pnasnexus/pgad036

44 Kerrigan, J. R., Veldhuis, J. D., Leyo, S. A., Iranmanesh, A. & Rogol, A. D. Estimation of daily cortisol production and clearance rates in normal pubertal males by deconvolution analysis. J Clin Endocrinol Metab 76, 1505–1510 (1993). 10.1210/jcem.76.6.8501158

45 Løvås, K. & Husebye, E. S. Continuous subcutaneous hydrocortisone infusion in Addison’s disease. Eur J Endocrinol 157, 109–112 (2007). 10.1530/eje-07-0052

46 Pang, Z. et al. MetaboAnalyst 6.0: towards a unified platform for metabolomics data processing, analysis and interpretation. Nucleic Acids Research 52, W398–W406 (2024). 10.1093/nar/gkae253

47 Carvalho, B. S. & Irizarry, R. A. A framework for oligonucleotide microarray preprocessing. Bioinformatics 26, 2363–2367 (2010). 10.1093/bioinformatics/btq431

48 Ritchie, M. E. et al. limma powers differential expression analyses for RNA-sequencing and microarray studies. Nucleic Acids Research 43, e47–e47 (2015). 10.1093/nar/gkv007

49 Rohart, F., Gautier, B., Singh, A. & Le Cao, K. A. mixOmics: An R package for ‘omics feature selection and multiple data integration. PLoS Comput Biol 13, e1005752 (2017). 10.1371/journal.pcbi.1005752

50 Bick, C., Gross, E., Harrington, H. A. & Schaub, M. T. What Are Higher-Order Networks? SIAM Review 65, 686–731 (2023). 10.1137/21m1414024

51 Masoomy, H., Askari, B., Tajik, S., Rizi, A. K. & Jafari, G. R. Topological analysis of interaction patterns in cancer-specific gene regulatory network: persistent homology approach. Sci Rep 11, 16414 (2021). 10.1038/s41598-021-94847-5

