## Supplementary Figures and Methods for "Higher-Order Organization Links Circulating Metabolites to Glucocorticoid Action in Human Adipose Tissue"

### SUPPLEMENTARY DATA

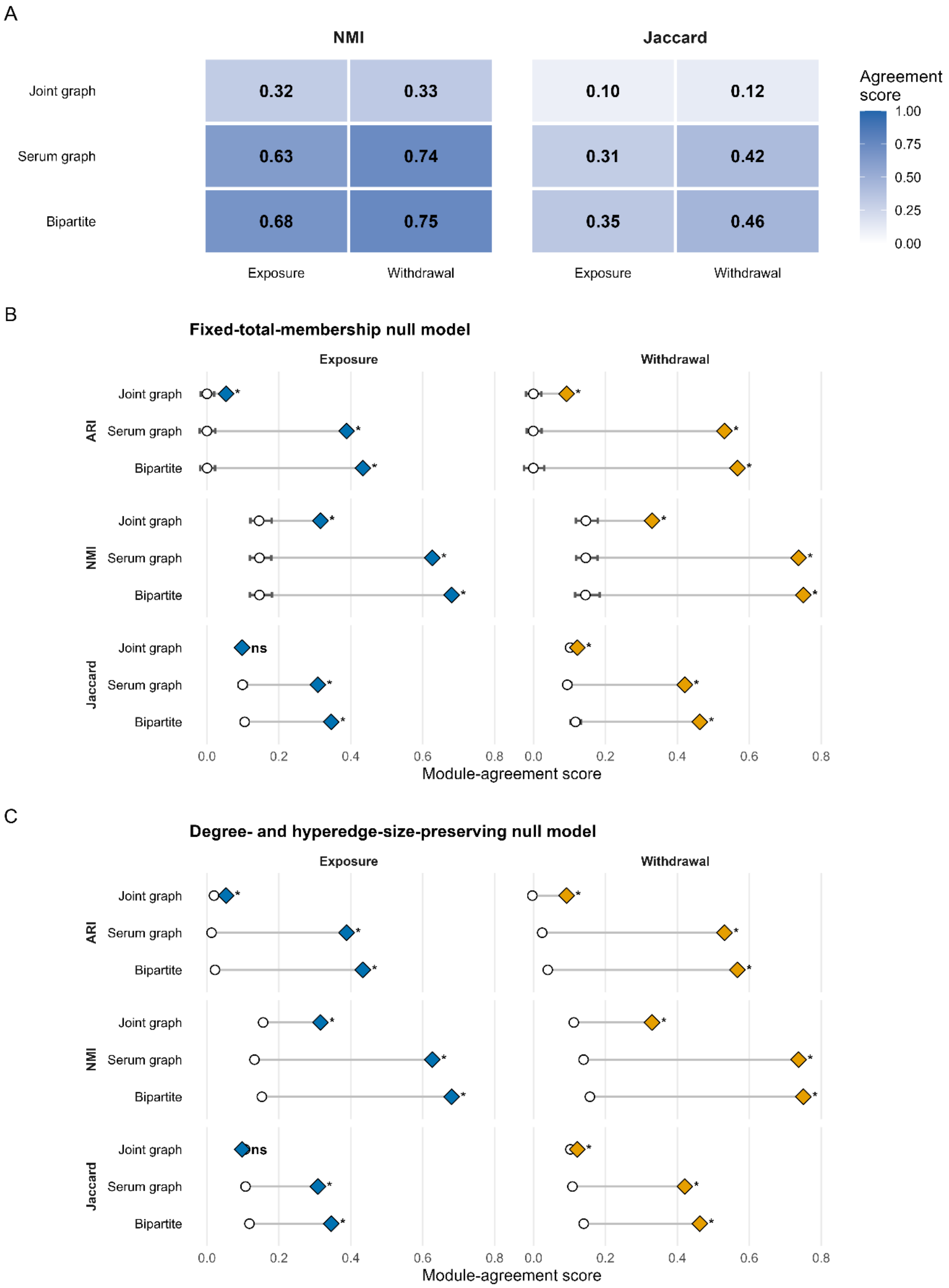

***Supplementary Figure 1. Alternative measures of module similarity between pairwise and higher-order network representations. (A)*** *Heatmaps showing normalised mutual information (NMI) and Jaccard similarity between serum metabolite modules derived from bipartite, serum correlation graph, and joint serum-adipose correlation graph approaches relative to hypergraph-derived modules under Exposure and Withdrawal conditions. NMI quantifies similarity in clustering information content, whereas the Jaccard index quantifies overlap among co-clustered metabolite pairs. Higher values indicate greater concordance between module assignments.* ***(B-C)*** *Observed Adjusted Rand Index (ARI), NMI and Jaccard overlap compared with null expectations generated using a* ***(B)*** *fixed-total-membership randomisation and* ***(C)*** *degree- and hyperedge-size preserving randomisation. Grey horizontal intervals indicate the central 95% of agreement scores across 1,000 null hypergraphs, white circles indicate null medians, and coloured diamonds indicate observed values. Asterisks indicate FDR-adjusted empirical greater-tail q-values; * q < 0.05; ns, not significant.*

***Supplementary Table 1. Metabolite classes of top loop-enriched metabolites in GC exposure and withdrawal networks.*** *The table lists metabolite classes corresponding to metabolites that appeared in at least ten more persistent loops in either the exposure or withdrawal network (top hits from Extended Data Fig. 2c)*

| **Functional class** | **More H1 features in Exposure ($\Delta \geq 10$)** | **More H1 features in Withdrawal ($\Delta \leq -10$)** |
|---|---|---|
| **Acylcarnitines** | Octanoylcarnitine | Myristoylcarnitine; Stearoylcarnitine |
| **Fatty acids (free)** | — | Oleic acid; Linoleic acid; Arachidonic acid; Palmitic acid; Octadecenoic acid |
| **Phospholipids (PC species)** | PC(P-16:0/0:0) or PC(0-16:1/0:0) | PC(18:3/0:0) linolenoyl; PC(16:0/0:0) palmitoyl; PC(17:0/0:0) |
| **Bile acids** | Taurodeoxycholate | Chenodeoxycholic acid glycine conjugate; Glycodeoxycholate |
| **Amino acids & derivatives** | L-Tryptophan; Gamma-Glu-Leu; 2-Oxoisocaproic acid; Pyroglutamic acid; | L-Kynurenine; Phenylalanine; Citrulline |
| **Neurotransmitters / xanthines** | Norepinephrine (noradrenaline) | Theobromine |
| **Organic acids (TCA / related)** | Alpha-ketoglutaric acid; Malic acid | Alanine |
| **Sugars & polyols** | Maltose; myo-Inositol | scyllo-Inositol |
| **Sterols** | Cholesterol | — |
| **Nucleotides / bases** | Adenosine-5-monophosphate; Uracil | — |
| **Phospho-headgroups / cofactors** | O-Phosphoethanolamine | — |

***Supplementary Table 2:*** *Serum metabolite bivariate predictive markers between GC exposure and withdrawal at 7am (second intervention day). Top 15 metabolite pairs, showing AUC, P-value and FDR.*
*Abbreviations: AUC, area under curve; FDR, false-discovery rate, LCMS, liquid chromatography mass spectrometry; GCMS, gas chromatography mass spectrometry*

| Metabolite Marker Pairs | AUC | P-value | FDR |
|---|---|---|---|
| Lactic Acid/Tryptophan | 1 | $3.62x10^{-5}$ | 0.040 |
| Heptadecanoic Acid/Tryptophan | 1 | $3.17x10^{-6}$ | 0.031 |
| S-Inositol/Kynurenine (GCMS) | 0.99 | $3.66x10^{-5}$ | 0.040 |
| Phenylalanine/Kynurenine | 0.98 | $7.58x10^{-5}$ | 0.050 |
| Kynurenine (LCMS)/S-Inositol | 0.98 | $1.15x10^{-5}$ | 0.038 |
| Kynurenine (LCMS)/Cholesterol | 0.98 | $2.11x10^{-5}$ | 0.040 |
| Cholesterol/Kynurenine (GCMS) | 0.98 | $3.58x10^{-5}$ | 0.040 |
| Kynurenine/Dodecanoic Acid | 0.97 | $7.32x10^{-6}$ | 0.036 |
| Kynurenine/Hexadecanoic Acid | 0.96 | $9.55x10^{-5}$ | 0.050 |
| Kynurenine/Heptadecanoic Acid | 0.96 | $5.04x10^{-5}$ | 0.045 |
| Tryptophan/Cholesterol | 0.96 | $6.01x10^{-5}$ | 0.049 |
| Kynurenine/1-Monostearoylglycerol | 0.95 | $8.87x10^{-5}$ | 0.050 |
| Pyroglutamic acid/Tryptophan | 0.95 | $2.93x10^{-5}$ | 0.040 |
| Hexadecanoic/Tryptophan | 0.95 | $4.79x10^{-5}$ | 0.045 |
| Tryptophan/Eicosatetraenoic Acid | 0.95 | $8.00x10^{-5}$ | 0.050 |

***Supplementary Table 3. GC-state dependent network connectivity of candidate serum metabolite pairs within GC-responsive adipose tissue features.*** *Pairwise co-degree represents the number of GC-responsive adipose metabolites and transcripts jointly associated with both serum metabolites forming each candidate ratio. Co-degree values were standardised separately within GC exposure and GC withdrawal relative to the distribution across the 15 evaluated candidate pairs. Positive Z-scores indicate greater shared adipose connectivity than the candidate-pair mean within that GC state.*

| Metabolite Pairs | Co-degree (Exposure) | Z-score (Exposure) | Co-degree (Withdrawal) | Z-score (Withdrawal) |
|---|---|---|---|---|
| Phenylalanine / Kynurenine | 24 | 2.92 | 11 | 0.0787 |
| Pyroglutamic acid / Tryptophan | 15 | 0.868 | 10 | -0.157 |
| Lactic acid / Tryptophan | 14 | 0.639 | 8 | -0.629 |
| Kynurenine (LCMS) / S-Inositol | 12 | 0.183 | 23 | 2.91 |
| Kynurenine (LCMS) / Cholesterol | 10 | -0.274 | 10 | -0.157 |
| Kynurenine / Dodecanoic acid | 12 | 0.183 | 11 | 0.0787 |
| Kynurenine / 1-Monostearoylglycerol | 12 | 0.183 | 6 | -1.10 |
| Kynurenine / Hexadecanoic acid | 11 | -0.0457 | 12 | 0.315 |
| Tryptophan / Cholesterol | 10 | -0.274 | 7 | -0.865 |
| Tryptophan / Eicosatetraenoic acid | 10 | -0.274 | 8 | -0.629 |
| S-Inositol / Kynurenine (GCMS) | 9 | -0.502 | 16 | 1.26 |
| Hexadecanoic acid / Tryptophan | 9 | -0.502 | 7 | -0.865 |
| Kynurenine / Heptadecanoic acid | 8 | -0.731 | 12 | 0.315 |
| Cholesterol / Kynurenine (GCMS) | 7 | -0.959 | 9 | -0.393 |
| Heptadecanoic acid / Tryptophan | 5 | -1.42 | 10 | -0.157 |

***Supplementary Table 4.*** *Top marker ratios and individual metabolites as markers predicting GC exposure from withdrawal at midnight time point in BIOCORT study.*

| Metabolite Marker Pair | AUC | P-value | FDR |
|---|---|---|---|
| Phenylalanine/Kynurenine | 0.90 | 0.000382 | 0.00420 |
| Tryptophan | 0.83 | 0.00788 | 0.0355 |
| Kynurenine | 0.82 | 0.00967 | 0.0355 |
| Pyroglutamic Acid/Tryptophan | 0.82 | 0.0179 | 0.0493 |
| Tryptophan/Eicosatetraenoic Acid | 0.75 | 0.117 | 0.257 |
| Tryptophan/Hexadecanoic Acid | 0.69 | 0.234 | 0.318 |
| Kynurenine/Hexadecanoic Acid | 0.67 | 0.173 | 0.318 |
| Pyroglutamic Acid | 0.63 | 0.408 | 0.499 |
| Eicosatetraenoic Acid | 0.63 | 0.391 | 0.537 |
| Hexadecanoic Acid | 0.59 | 0.537 | 0.591 |
| Phenylalanine | 0.56 | 0.591 | 0.591 |

***Supplementary Table 5*** *– Validation of the top marker pairs in dose-response study, paired t-test comparing high vs low GC dose after both acute (4 hours) and chronic (7 days) exposure. Where metabolites were found in both LCMS and GCMS, all combinations are reported. Table is sorted by chronic p-value.*

| Metabolite Marker pair | Acute P-value | Acute FDR | Chronic P-value | Chronic FDR |
|---|---|---|---|---|
| Phenylalanine (LCMS)/Kynurenine | 0.24 | 0.93 | 0.0045 | 0.082 |
| Phenylalanine (GCMS)/Kynurenine | 0.040 | 0.72 | 0.011 | 0.10 |
| Kynurenine/Heptadecanoic Acid | 0.37 | 0.93 | 0.017 | 0.10 |
| Kynurenine/s-Inositol | 0.26 | 0.93 | 0.024 | 0.11 |
| Kynurenine/Dodecanoic Acid | 0.48 | 0.93 | 0.031 | 0.11 |
| Tryptophan (GCMS)/Cholesterol | 0.92 | 0.93 | 0.052 | 0.16 |
| Heptadecanoic Acid/Tryptophan (GCMS) | 0.72 | 0.93 | 0.13 | 0.29 |
| Pyroglutamic Acid (GCMS)/Tryptophan (GCMS) | 0.50 | 0.93 | 0.13 | 0.29 |
| Tryptophan (GCMS)/Eicosatetraenoic Acid | 0.93 | 0.93 | 0.14 | 0.29 |

| | | | | |
|---|---|---|---|---|
| Lactic Acid/Tryptophan (GCMS) | 0.83 | 0.93 | 0.18 | 0.33 |
| Pyroglutamic Acid (LCMS)/Tryptophan (LCMS) | 0.63 | 0.93 | 0.21 | 0.33 |
| Tryptophan (LCMS)/Cholesterol | 0.74 | 0.93 | 0.22 | 0.33 |
| Pyroglutamic Acid (LCMS)/Tryptophan (GCMS) | 0.70 | 0.93 | 0.27 | 0.37 |
| Kynurenine/Cholesterol | 0.92 | 0.93 | 0.39 | 0.45 |
| Heptadecanoic Acid/Tryptophan (LCMS) | 0.74 | 0.93 | 0.64 | 0.73 |
| Pyroglutamic Acid (GCMS)/Tryptophan (LCMS) | 0.11 | 0.93 | 0.65 | 0.73 |
| Tryptophan (LCMS)/Eicosatetraenoic Acid | 0.40 | 0.93 | 0.79 | 0.84 |
| Lactic Acid/Tryptophan (LCMS) | 0.90 | 0.93 | 0.89 | 0.80 |

### SUPPLEMENTARY METHODS

**Adipose Tissue Sampling**

Subcutaneous needle biopsies were taken from the abdomen's right lower quadrant during saline infusion and from the left lower quadrant during hydrocortisone (HC) infusion. Following collection, excess blood was removed from the biopsies, the samples were promptly transferred to capped cryovials and then snap-frozen in liquid nitrogen.

Following the adipose tissue sampling, three microdialysis catheters were inserted into the abdominal subcutaneous fat in the abdomen's upper quadrants for interstitial metabolite profiling (adipose tissue metabolomics). The catheters were inserted in the abdomen's left upper quadrant during saline infusion and in the right upper quadrant during hydrocortisone (HC) infusion (opposite side where the adipose tissue biopsies were previously collected). The catheters (Asahi, Asahi Kasei Medical Co., Japan) were composed of polyethylene membrane with an inner diameter (ID) of 340 µm, an outer diameter (OD) of 440 µm, a pore size of 0.3 µm, and a molecular weight cut-off of 3 MDa. A needle attached at the outlet facilitated membrane insertion into the tissue. A borosilicate glass capillary tube (100 µL; A. Hartenstein GmbH, Würzburg, Germany) was positioned at approximately a 40° angle at the catheter outlet for sample collection. The catheter inlet was connected via tubing to a syringe placed in a CMA 100 microinjection pump (CMA Microdialysis AB, Kista, Sweden), and perfusion was maintained at a flow rate of 1.5 µL/min using isotonic saline as the perfusion fluid (PMID: 33712379). Following the sample collection that took approximately 2 hours, the collected dialysate was promptly transferred to capped cryovials. All specimens during the study interventions were stored at −80 °C until analysis. All analyses were performed in one run.

**LCMS Adipose Microdialysis Metabolomic Profiling**

LCMS was performed at the Swedish Metabolomics Center in Umeå, Sweden. Prior to sample extraction 300µL of "microdialysis fluid" (dialysates) were freeze-dried overnight. 220µL of extraction buffer (80/20 v/v methanol: water) including internal standards for LCMS (see supplementary) were added to the tubes. The sample was shaken at 30 Hz for 2 minutes in a mixer mill. The sample was centrifuged at +4 °C, 14 000 rpm, for 10 minutes. The supernatant, 200µL was transferred to micro vials and evaporated to dryness under a stream of nitrogen gas, the samples were stored at -80 °C until analysis.

Before LCMS analysis the sample was re-suspended in 20 + 20 µL methanol and water. The samples were first analyzed in positive mode. After all samples had been analyzed, the instrument was switched to negative mode and a second injection of each sample was performed.

The chromatographic separation was performed on an Agilent 1290 Infinity UHPLC-system (Agilent Technologies, Waldbronn, Germany). 5 µL of each sample were injected onto an Acquity UPLC HSS T3, 2.1 x 50 mm, 1.8 µm C18 column in combination with a 2.1 mm x 5 mm, 1.8 µm VanGuard precolumn (Waters Corporation, Milford, MA, USA) held at 40 °C. The gradient elution buffers were A (H2O, 0.1 % formic acid) and B (75/25 acetonitrile:2-propanol, 0.1 % formic acid), and the flow-rate was 0.5 mL min-1. The compounds were eluted with a linear gradient consisting of 0.1 - 10 % B over 2 minutes, B was increased to 99 % over 5 minutes and held at 99 % for 2 minutes; B was decreased to 0.1 % for 0.3 minutes and the flow-rate was increased to 0.8 mL min-1 for 0.5 minutes; these conditions were held for 0.9 minutes, after which the flow-rate was reduced to 0.5 mL min-1 for 0.1 minutes before the next injection.

The compounds were detected with an Agilent 6546 Q-TOF mass spectrometer equipped with a jet stream electrospray ion source operating in positive or negative ion mode. The settings were kept identical between the modes, with the exception of the capillary voltage. A reference interface was connected for accurate mass measurements; the reference ions purine (4 µM) and HP-0921 (Hexakis(1H, 1H, 3H-tetrafluoropropoxy)phosphazine) (1 µM) were infused directly into the MS at a flow rate of 0.05 mL min-1 for internal calibration, and the monitored ions were purine m/z 121.05 and m/z 119.03632; HP-0921 m/z 922.0098 and m/z 966.000725 for positive and negative mode respectively. The gas temperature was set to 150°C, the drying gas flow to 8 L min-1 and the nebulizer pressure 35 psig. The sheath gas temp was set to 350°C and the sheath gas flow 11 L min-1. The capillary voltage was set to 4000 V in positive ion mode, and to 4000 V in negative ion mode. The nozzle voltage was 300 V. The fragmentor voltage was 120 V, the skimmer 65 V and the OCT 1 RF Vpp 750 V. The collision energy was set to 0 V. The m/z range was 70 - 1700, and data was collected in centroid mode with an acquisition rate of 4 scans s-1 (1977 transients/spectrum).

**Hypergraphs**

A hypergraph $H = (N,E)$ consists of a collection of nodes ($N$) and hyperedges ($E$), where each hyperedge can connect more than two nodes simultaneously. In this study, definitions of nodes and hyperedges depended on the specific analysis.

Associations between nodes and hyperedges were quantified using Pearson correlation coefficients, calculated across samples within each GC state. Binary node-hyperedge membership was defined by thresholding at the standard deviation of the condition-specific correlation matrix. Correlations with an absolute value greater than this threshold were assigned a value of 1, and all remaining entries were given a value of 0, yielding a binary incidence matrix $H \in \{0,1\}^{N \times E}$(with $N$ nodes and $E$ hyperedges) which defines the full hypergraph. Each row corresponds to a node, and each column a hyperedge. Unless otherwise stated, thresholding was applied within the GC-state-specific correlation matrix. For Δ shared-connectivity comparisons, the withdrawal incidence matrix was thresholded using the exposure-derived correlation standard deviation to preserve comparability across conditions.

**Pairwise versus Hypergraph Module Analysis**

Three alternative network representations were constructed for comparison with the serum-adipose hypergraph. The serum-correlation graph was generated from pairwise correlations among serum metabolites and represents conventional metabolite-network organisation. The joint serum-adipose correlation graph was generated by combining serum metabolites, adipose metabolites and adipose transcripts into a single feature matrix, followed by calculation of all pairwise feature-feature correlations. Module assignments corresponding to serum metabolites were subsequently extracted for comparison with hypergraph-derived modules. The bipartite incidence representation was derived directly from the serum-adipose incidence matrix and preserves direct serum-adipose associations without introducing higher-order relationships between serum metabolites.

For the hypergraph representation, a projected adjacency matrix was constructed from the incidence matrix and hierarchical clustering was performed on the resulting similarity matrix. For serum and joint-correlation graphs, pairwise Pearson-correlation matrices were calculated following row-wise standardisation, thresholded using a standard-deviation-based correlation threshold, converted to similarity matrices, and clustered using agglomerative hierarchical clustering (average linkage). For bipartite analyses, serum metabolites were clustered directly from binary incidence profiles using binary distance and hierarchical clustering. Module assignments were generated using a fixed cut of $k = 15$ clusters.

These alternative representations were selected because they capture distinct aspects of the underlying data. The serum graph represents conventional pairwise organisation among circulating metabolites, the joint graph represents pairwise structure across the full multi-omic dataset, and the bipartite representation preserves direct serum-adipose relationships while excluding higher-order metabolite interactions. Comparison with the hypergraph therefore enabled assessment of whether observed module organisation could be explained by conventional pairwise structure or reflected higher-order organisation encoded within the hypergraph.

**Hypergraph Null Models**

To assess whether observed hypergraph organisation exceeded that expected by chance, two complementary null-model frameworks were applied independently to the exposure and withdrawal incidence matrices. For each condition, 1,000 null hypergraphs were generated

The fixed-membership null model preserved the total number of node-hyperedge memberships while randomising their allocation within the incidence matrix. For each randomisation, the observed number of memberships was distributed uniformly across the incidence matrix by randomly selecting matrix positions without replacement and assigning them a value of one, while all remaining entries were assigned zero. This approach preserves overall hypergraph density but does not preserve node hyperdegree or hyperedge-size distributions.

A second null model was generated using a Curveball-style randomisation procedure. Starting from the observed binary incidence matrix, repeated trades of hyperedge memberships were performed between randomly selected node pairs[1]. This procedure preserves both node hyperdegree (row sums) and hyperedge size (column sums) while randomising higher-order organisation. The target number of effective trades was set to twenty times the number of nodes in the incidence matrix. Preservation of row and column sums was verified for every randomised hypergraph.

For module analyses, modules were recalculated for each null hypergraph using the same hierarchical-clustering procedure applied to the observed hypergraph. Module similarity metrics (ARI, NMI, Jaccard overlap, and pairwise agreement) were then compared with null distributions to calculate empirical P-values and FDR-adjusted P-Value.

For structural analyses, node hyperdegree, hyperedge size, shared connectivity and the corresponding entropy metrics were recalculated for every null hypergraph. Two entropy measures were calculated from the projected co-degree matrix. Pairwise co-degree entropy quantified how evenly shared hyperedge memberships were distributed across serum-metabolite pairs. Summed co-degree entropy quantified how evenly total shared connectivity was distributed across serum metabolites. Lower values indicate that pairwise co-degree or summed co-degree is concentrated among a smaller subset of metabolite pairs or metabolites, respectively. Observed values were compared with the corresponding null distributions using empirical P-values and Z-scores.

**Shared-Connectivity Analysis**

To assess shared participation of serum metabolites in adipose-linked associations, we derived connectivity measures. Each element of the incidence matrix indicates whether a given serum metabolite is associated with a specific adipose feature based on the defined correlation threshold. For each metabolite, node hyperdegree was defined as the number of adipose features with which that metabolite was associated (i.e. the row sum of the incidence matrix), representing direct pairwise serum–adipose associations.

To summarise higher-order coordination, a reduced adjacency matrix was constructed as the product of the incidence matrix and its transpose $A = HH^T$. Each entry of this matrix represents the number of adipose features jointly associated with a pair of serum metabolites, which is their pairwise co-degree. This is consistent with definitions of co-degree as shared hyperedge membership in hypergraphs[2]. The diagonal elements of the adjacency matrix were set to zero prior to calculating shared connectivity to remove self-counting of node degree. For each metabolite, shared connectivity was calculated as the row sum of the off-diagonal co-degree matrix, equivalent to a summed co-degree across all other metabolites.

To compare these quantities across conditions, both node degree and shared connectivity were min–max normalised within each GC state, and a summary metric (Δ shared-connectivity score) was defined as the difference between shared connectivity and node degree.

**Marker-Pair Enrichment Analysis**

For each GC state, higher-order overlap among candidate serum marker metabolites was quantified using pairwise co-degree, defined as the number of adipose features jointly associated with both metabolites. This corresponds to the entries of the reduced adjacency matrix as described above.

For each pair of serum metabolites $a$ and $b$, pairwise co-degree was defined as:

$$c_{ab} = \sum_{k=1}^{m} H_{ak} H_{bk}$$

where $H_{ak}$ and $H_{bk}$ indicate membership of metabolites $a$ and $b$ in hyperedge $k$, and $m$ is the total number of adipose features included in the restricted incidence matrix.

This formulation represents the pairwise co-degree between metabolites, capturing the extent to which candidate serum markers share adipose-linked association patterns. Shared connectivity was computed separately for GC exposure and GC withdrawal. To enable comparison across metabolite pairs within each condition, shared connectivity values were standardised using Z-scores.

**Pathway Entropy Analysis**

Metabolic pathways were defined using Gene Ontology gene sets, including amino acid metabolism, carbohydrate metabolism, lipid metabolism, nucleobase metabolism, and proteolysis.

For each pathway, pathway genes were first matched to adipose transcriptomic expression matrices under each GC state. Where transcripts contained multiple names, rows were retained if any name matched a pathway gene. Because pathway sizes varied, a fixed number of 50 genes were randomly sampled without replacement from each iteration. This resampling procedure was repeated 300 times per pathway per condition, generating condition-specific distributions of entropy estimates while controlling for pathway-size effects.

For each iteration, the sampled pathway genes were correlated against the remainder of the adipose transcriptome. The resulting correlation was converted to absolute values, thresholded at its standard deviation, and binarised to produce a pathway-specific incidence matrix. A reduced adjacency matrix was then derived, capturing the number of shared associations between pairs of sampled pathway genes. Shannon entropy was then computed from the resulting adjacency matrix values using the entropy package in R, providing a summary of how evenly higher-order shared associations were distributed across pathway gene pairs.

To control for global transcriptome-wide changes, random gene sets of identical size (n = 50) were sampled and analysed using the same procedure, providing a negative control for non-specific changes in network organisation.

Differences in entropy between GC exposure and withdrawal were assessed using Bayesian two-sample t-tests implemented in the BayesFactor package[3]. For each pathway, posterior samples of the group-difference parameter were drawn from the fitted model (10,000 posterior iterations), and highest-density intervals were used to visualise uncertainty in the direction and magnitude of entropy change.

**Topological Data Analysis**

**Distance Matrix Construction**

TDA was employed to capture structural and connectivity patterns in the serum-adipose hypergraph. The binary incidence matrix was first converted into a pairwise distance matrix between serum metabolites.

For each pair of metabolites, the Jaccard index was computed from the hyperedge membership profiles:

$$J(i,j) = \frac{|H_i \cap H_j|}{|H_i \cup H_j|}$$

where $H_i$ and $H_j$ denote the hyperedge memberships of metabolites $i$ and $j$ .

Jaccard distance was then calculated as:

$$d_J(i,j) = 1 - J(i,j)$$

Smaller values indicate more similar hyperedge profiles and larger distances indicate distinct profiles. The resulting $n \times n$ distance matrix, where $n$ is the number of serum metabolites, was used as the direct input for the Python persistent homology workflow.

**Persistent Homology Computation**

Persistent homology for persistence diagrams, barcode visualisation, and Wasserstein comparisons were computed in Python using Ripser[4], where homology was calculated up to the second dimension. The distance matrix is treated as a point cloud embedded in abstract metric space. The Vietoris-Rips filtration grows balls of increasing radius around each point, and simplicial complexes are constructed at each scale parameter $\varepsilon$. The emergence and disappearance of topological features were tracked across each filtration to identify:

- $H_0$ – connected components
- $H_1$ – loops
- $H_2$ – voids

Persistence diagrams were generated for the $0^{th}$, $1^{st}$, and $2^{nd}$ homology dimensions. To compare persistence diagrams, Wasserstein (earth-mover) distances were computed using the Persim library. This metric quantifies the minimal cost of transforming one diagram into another, providing a measure of topological similarity.

**Subsampling and permutation testing**

To assess the robustness of topological differences to node composition, node-subsampled pseudo-replicate persistence diagrams were generated from each condition-specific distance matrix. For each GC state, 200 pseudo-replicates were created by randomly selecting 80% of nodes without replacement, extracting the induced distance submatrix, and recomputing persistent homology for H0-H2.

For each homology dimension (H0, H1, H2) pairwise Wasserstein distances between all pseudo-replicate diagrams were computed using persim and classified as intra-condition (exposure-exposure or withdrawal-withdrawal), or inter-condition (exposure-withdrawal). Condition separation was summarised as the difference between the median inter-condition distance and the median intra-condition distance. Statistical significance was assessed using a label-permutation test in which pseudo-replicate condition labels were shuffled while preserving group sizes, and the median(inter-condition) – median(intra-condition) statistic was recalculated for each permutation (n=5000)[5].

For persistence-summary analyses, each pseudo-replicate diagram was additionally summarised by the number of finite features, total persistence, median persistence, and maximum persistence in each homology dimension. Prior to these summary calculations, infinite points were removed and a minimum persistence threshold of 0.01 was applied, to remove trivial features from calculations. Exposure and withdrawal distributions for each metric were compared using label-permutation tests on the difference in group medians.

**Representative cycle extraction**

Persistent homology analyses were additionally performed in Julia v1.7.3 using the *Eirene* package. This package was used specifically because it computes matroid generators for homology classes, allowing extraction of representative cycles for H1 features. This provides a principled and reproducible identification of metabolites participating in each persistent loop, enabling biological interpretation of higher-order structures[6]. Condition-specific pairwise and Jaccard distance matrices were imported with metabolite labels. Persistent homology was computed using the Vietoris–Rips filtration up to homology dimension 2 for both GC exposure and GC withdrawal. Barcode tables for H0, H1, and H2 were exported with class index, birth, death, and persistence values. Representative H1 cycle edge lists were also exported for each barcode class, including metabolite-name annotations. These Eirene-derived outputs were used for downstream comparison and visualisation of persistent loop structures.

**H1 Loop Analysis**

To characterise metabolite participation in persistent H1 loops, barcode class tables and H1 class-representative files exported from the Eirene workflow were analysed. H1 barcode classes with persistence ≥ 0.01 were retained for loop-participation analysis. For each retained class representative, metabolites were counted once per loop if they appeared in that loop structure. Condition-specific loop-participation counts were then compared, and metabolites showing an absolute difference of at least 10 loop appearances between GC exposure and GC withdrawal were highlighted and grouped by functional class for visualisation.

**Cosinor Analysis**

In the DUOCORT study, metabolite and cortisol values were first z-scored within individuals to standardise scale across participants and analytes. Group-level mean z-scored values and standard errors of the mean (SEM) were then calculated at each time point. Circadian rhythmicity was modelled using a single-component 24-hour cosinor formulation implemented as a linear model of the form:

$$y(t) = M + \beta_c \cos(\omega t) + \beta_s \sin(\omega t)$$

where:

$$\omega = \frac{2\pi}{24x60}$$

for time expressed in minutes. Separate group-level models were fitted for each metabolite and dosing regimen (once-daily and thrice-daily), and fitted curves were used for visualisation alongside observed means ± SEM. P-values displayed on the cosinor panels correspond to the fitted group-level model for the relevant analyte and dosing regimens.

#### SUPPLEMENTAL REFERENCES


1 Chodrow, P. S. Configuration models of random hypergraphs. *Journal of Complex Networks* **8** (2020). https://doi.org:10.1093/comnet/cnaa018

2 Jiang, T. & Ma, J. Cycles of given lengths in hypergraphs. *Journal of Combinatorial Theory, Series B* **133**, 54-77 (2018). https://doi.org/10.1016/j.jctb.2018.04.004

3 Rouder, J. N., Speckman, P. L., Sun, D., Morey, R. D. & Iverson, G. Bayesian t tests for accepting and rejecting the null hypothesis. *Psychonomic Bulletin & Review* **16**, 225-237 (2009). https://doi.org:10.3758/PBR.16.2.225

4 Tralie, C., Saul, N. & Bar-On, R. Ripser.py: A Lean Persistent Homology Library for Python. *Journal of Open Source Software* **3** (2018). https://doi.org:10.21105/joss.00925

5 Robinson, A. & Turner, K. Hypothesis testing for topological data analysis. *Journal of Applied and Computational Topology* **1**, 241-261 (2017). https://doi.org:10.1007/s41468-017-0008-7

6 Henselman, G. & Ghrist, R. Matroid filtrations and computational persistent homology. *arXiv preprint arXiv:1606.00199* (2016).
